# Identify compound-protein interaction with knowledge graph embedding of perturbation transcriptomics

**DOI:** 10.1101/2024.04.08.588632

**Authors:** Shengkun Ni, Xiangtai Kong, Yingying Zhang, Zhengyang Chen, Zhaokun Wang, Zunyun Fu, Ruifeng Huo, Xiaochu Tong, Ning Qu, Xiaolong Wu, Kun Wang, Wei Zhang, Runze Zhang, Zimei Zhang, Jiangshan Shi, Yitian Wang, Ruirui Yang, Xutong Li, Sulin Zhang, Mingyue Zheng

## Abstract

The emergence of perturbation transcriptomics provides a new perspective and opportunity for drug discovery, but existing analysis methods suffer from inadequate performance and limited applicability. In this work, we present PertKGE, a method designed to improve compound-protein interaction with knowledge graph embedding of perturbation transcriptomics. PertKGE incorporates diverse regulatory elements and accounts for multi-level regulatory events within biological systems, leading to significant improvements compared to existing baselines in two critical “cold-start” settings: inferring binding targets for new compounds and conducting virtual ligand screening for new targets. We further demonstrate the pivotal role of incorporating multi- level regulatory events in alleviating dataset bias. Notably, it enables the identification of ectonucleotide pyrophosphatase/phosphodiesterase-1 as the target responsible for the unique anti-tumor immunotherapy effect of tankyrase inhibitor K-756, and the discovery of five novel hits targeting the emerging cancer therapeutic target, aldehyde dehydrogenase 1B1, with a remarkable hit rate of 10.2%. These findings highlight the potential of PertKGE to accelerate drug discovery by elucidating mechanisms of action and identifying novel therapeutic compounds.

## Introduction

Identifying compound-protein interaction (CPI) is fundamental for developing therapeutic compounds and understanding their target-level mechanisms of action (MOA)^1^. Over the past few decades, numerous in silico methods have been proposed and widely used in drug discovery pipelines due to their cost-effectiveness and high-throughput capabilities, offering valuable insights and guidance for both in vitro and in vivo experiments^2,3^.

In the past decades, a significant amount of biological data has been accumulated. This has led to computational methods evolving from primarily relying on cheminformatics and structural biology to incorporating multiple perspectives. For instance, systematic profiling of small-molecule perturbation effects, including cell images^4^, transcriptomics^5,6^, proteomics^7^ and metabolomics^8^, offers new opportunities to identify CPI^9,10^. Among these omics data, perturbation transcriptomics, which captures a ‘snapshot’ of differential mRNA abundance after perturbation, has received the most extensive attention due to its high-throughput nature and ability to acquire large-scale data^5,6,11,12^. However, inherent noise in biological data^13^, cellular homeostasis^14^, and dynamic changes in mRNA expression^15^, make CPI not directly reflected in the most differentially expressed genes (DEGs)^16^. This makes predicting CPI based on perturbation transcriptomics a challenging task.

Several methods can be used to address this problem, including comparative analysis and causal reasoning^17^. Comparative analysis involves finding the appropriate similarity between the query profile and well-annotated reference profiles, then assigning the MOA of the most similar reference profile to the query. A notable example is the Connectivity Map (CMap), which uses a connectivity score based on gene set enrichment analysis (GSEA) to measure the similarity^5^. Recently, some studies have introduced machine learning (ML)-based similarity to improve performance^18–20^. We also developed a deep-learning based method, SSGCN, to discover the hidden correlations between compound perturbation profiles and gene knockdown profiles^21^. However, these methods may not be directly applicable when analyzing CPI related to newly studied targets lacking relevant reference profiles.

Causal reasoning employs a systematic biological perspective, utilizing a prior knowledge network (PKN) to construct causal link and locate upstream nodes that can most accurately explain the observed downstream mRNA expression changes^17^. For example, DEMAND combines the gene regulatory network (GRN) and protein–protein interaction (PPI) to infer targets, assuming that compounds influence the expression of downstream genes^16^. ProTINA employs a dynamic model of cell-type-specific protein-gene transcriptional regulation to identify targets with high scores of network dysregulation^22^. A recent approach, FL-DTD, builds tissue-specific biological networks by integrating five preliminary networks and infers targets through a feedback loop assumption^23^. While these strategies have yielded effective tools for predicting CPI across any protein in the PKN, they face two main challenges. First, these methods tend to overlook key regulatory events, resulting in the omission of crucial regulatory patterns in the PKN. Second, these methods rely on known biology-inspired assumptions, which may not capture complex or as-yet-ununderstood expression patterns.

In recent years, knowledge graphs have become a promising method for integrating and analyzing multi-omics data^24^. Several curated biomedical knowledge graphs (BKGs), like HetioNet^25^, BioKG^26^, PharmKG^27^, and PrimeKG^28^, have been created for downstream analysis. However, directly analyzing high-dimensional transcriptomic data based on these BKGs is not appropriate. On the one hand, these BKGs contain a significant amount of redundant knowledge less related to chemical perturbation, such as diseases, side effects, anatomies, etc. On the other hand, the interactions between genes in these BKGs are too coarse-grained to finely represent the cellular response to chemical perturbation.

Here, we introduce PertKGE to improve CPI prediction based on perturbation transcriptomics by constructing biologically meaningful knowledge graph. Unlike other BKGs, this knowledge graph breaks down genes into DNAs, messenger RNAs (mRNAs), long non-coding RNAs (lncRNAs), microRNAs (miRNAs), transcription factors (TFs), RNA-binding proteins (RBPs) and other proteins. This enables PertKGE to consider various fine-grained interactions between genes to simulate post-transcriptional and post-translational regulatory events in biological system, which intuitively aligns more closely with real world cellular responses to chemical perturbations. Then, PertKGE uses the knowledge graph embedding (KGE) algorithm, DistMult^29^, to create knowledge-rich dense vectors and make CPI prediction based on the feature vectors. Compared to other baselines, PertKGE exhibited better performance in two cold-start settings while having a broader scope of application. We then conducted a comparison of our knowledge graph with other BKGs and performed an ablation study. The results showed that our knowledge graph enhanced the connections between genes and alleviated the impact of dataset bias on ML models. The ability of PertKGE in practical applications was also validated through biochemical experiments in this study. By combining PertKGE and experimental verifications, we successfully identified ectonucleotide pyrophosphatase/phosphodiesterase-1 (ENPP1) as the target influencing the immune phenotype for a tankyrase (TNKS) inhibitor K-756, and discovered five novel scaffold hits of aldehyde dehydrogenase 1B1 (ALDH1B1). These results demonstrate that PertKGE can be a valuable tool for predicting CPI from perturbation transcriptomics.

## Results

### Overview of PertKGE

The workflow of PertKGE can be divided into three parts: (1) construction of biologically meaningful knowledge graph; (2) train stage for obtaining the knowledge-rich embedding; (3) inference stage to give recommendation list.

### Construction of biologically meaningful knowledge graph

Drawing from causal reasoning^17^, we view a compound’s binding to one or more cellular targets as the cause, and the observed DEGs as the effect. This cause and effect are connected by a process involving various cellular regulatory events, which can be either linear or complex. Based on this concept, we construct a new knowledge graph with biological meaning by collecting ordered triples in the format of (head, relation, tail) from three components (Fig. 1A).

**Fig. 1.**
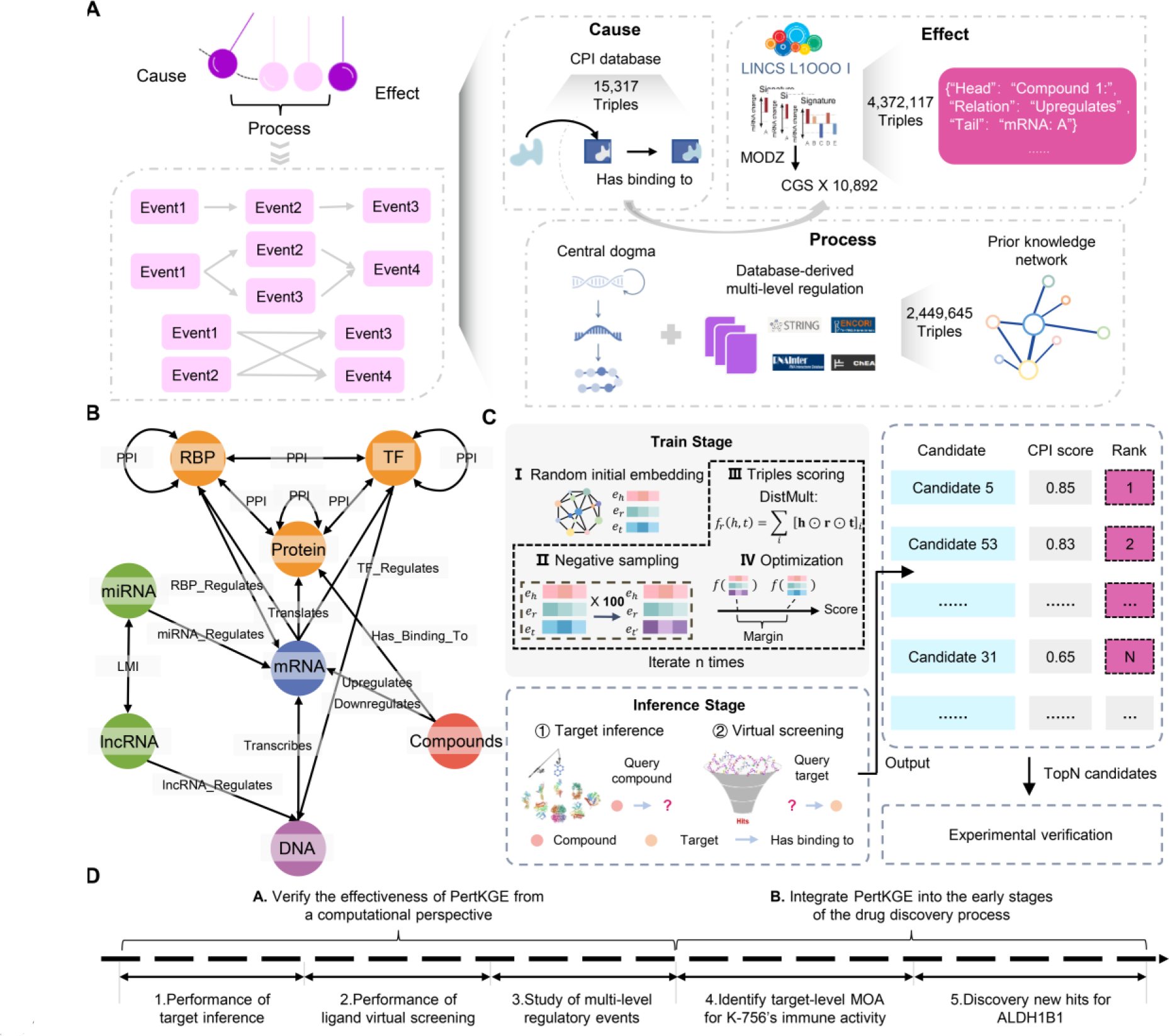
Overview of PertKGE. A, The construction pipeline of biologically meaningful knowledge graph. B, The graph schema of knowledge graph. The (Compound, Has_Binding_To, TF) and (Compound, Has_Binding_To, RBP) have been omitted due to their limited quantities. C, A schematic diagram illustrating the training and inference of PertKGE. D, Two stages demonstrating the effectiveness of PertKGE.

Effect component: This component leverages level 5 compound-induced transcriptomic data (known as signatures) from the Library of Integrated Network-Based Cellular Signatures (LINCS) Phase I^6^. Our previous work indicates that perturbations on PC-3 cells exhibit the best performance in CPI prediction^21^. Therefore, we only considered signatures from PC-3 cells. For each compound, moderated z-score (MODZ) was used to integrate multiple signatures obtained under different experimental conditions and generate a consensus gene signature (CGS)^6^. Consequently, we gathered 10,892 CGSs and processed the 200 most upregulated and downregulated genes from each, yielding a total of 4,372,117 triples like (Compound, Upregulated, mRNA) and (Compound, Downregulated, mRNA).

Cause component: This component comprises triples like (Compound, Has_Binding_To, Target) for 10,892 compounds, collected from multiple CPI databases^6,30–32^. Among these, only 2,845 compounds have a total of 15,317 CPI annotations, implying that 74% compounds lack CPI annotations.

Process component: This component leverages prior biological knowledge by incorporating essential regulatory events from various databases^33–36^ (see Method for more details). This component essentially captures the PKN encompassing 2,449,645 regulatory events rooted in the central dogma and multi-level regulatory elements, such as miRNAs, lncRNAs, RBPs, TFs and other proteins.

Finally, the three components were merged through entity alignment to form complete knowledge graph under the semantics of chemical perturbation. This knowledge graph can be represented as a directed heterogeneous graph, with nodes representing entities and edges representing relationships. Fig. 1B presents the graph schema of the knowledge graph and Table 1, 2 provide details about entities and relations (refer to Fig. S1 for more network analysis).

**Table 1.**
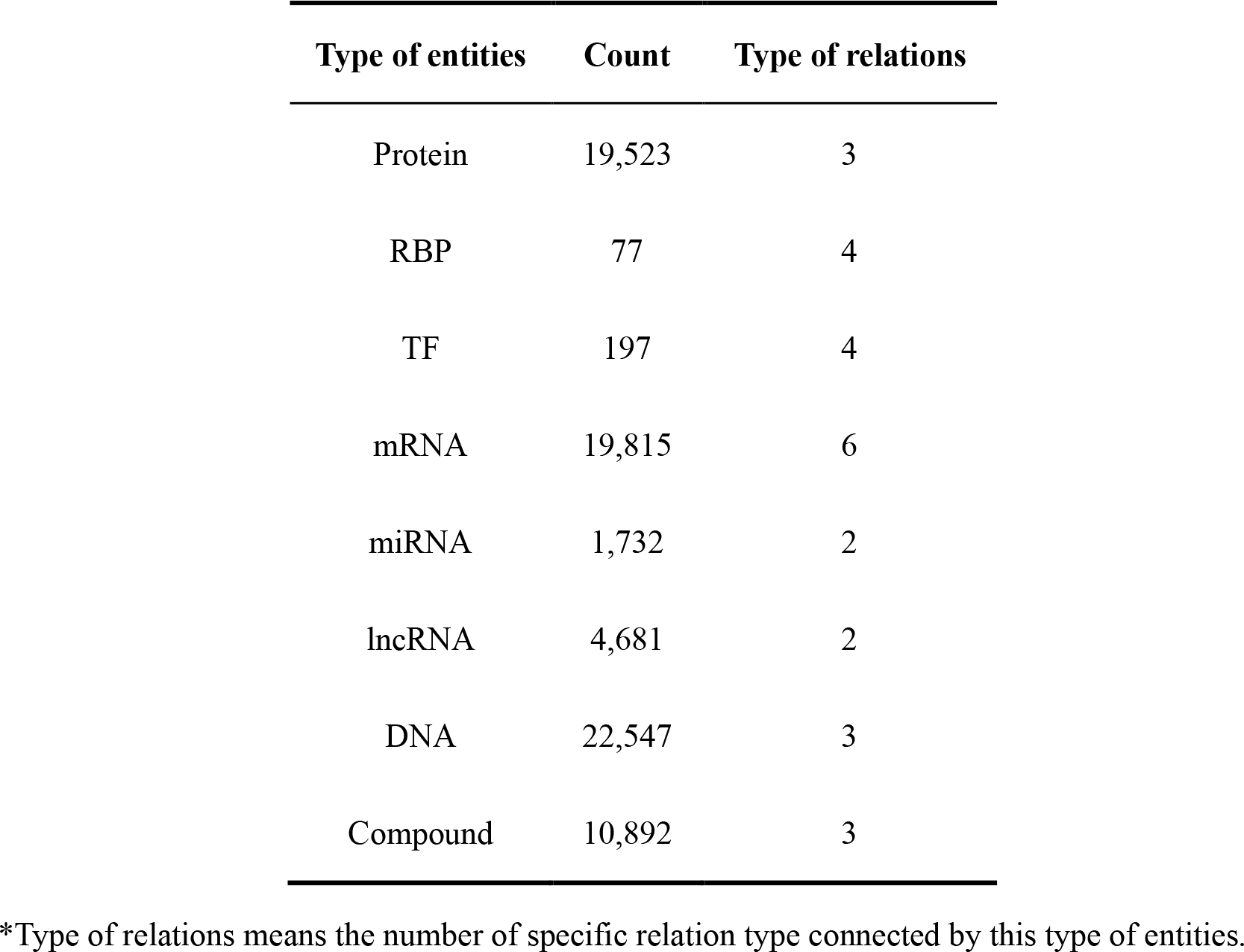
Entities in the chemical perturbation knowledge graph.

| Type of entities | Count | Type of relations |
| --- | --- | --- |
| Protein | 19,523 | 3 |
| RBP | 77 | 4 |
| TF | 197 | 4 |
| mRNA | 19,815 | 6 |
| miRNA | 1,732 | 2 |
| lncRNA | 4,681 | 2 |
| DNA | 22,547 | 3 |
| Compound | 10,892 | 3 |

\*Type of relations means the number of specific relation type connected by this type of entities.

**Table 2.**
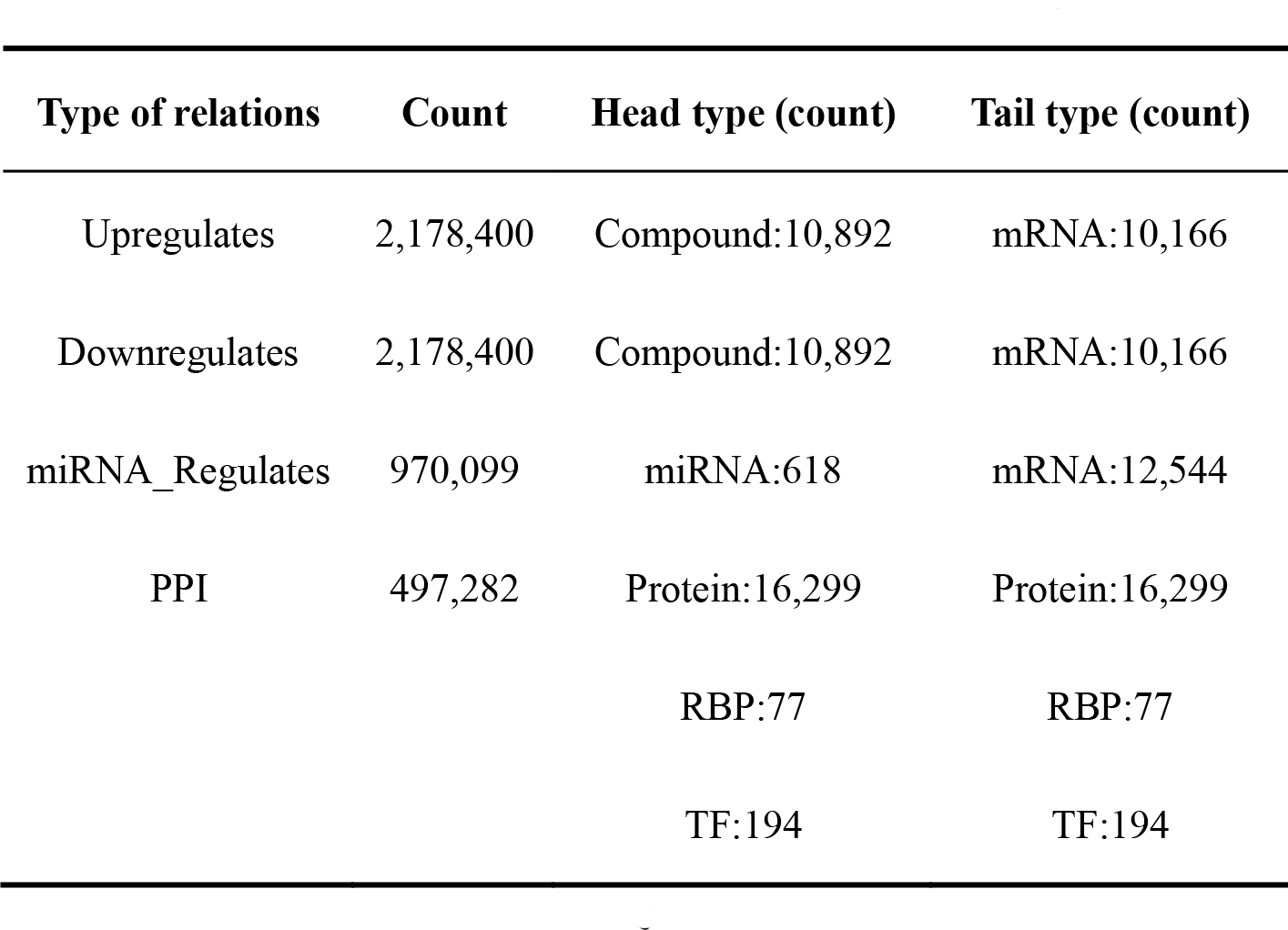
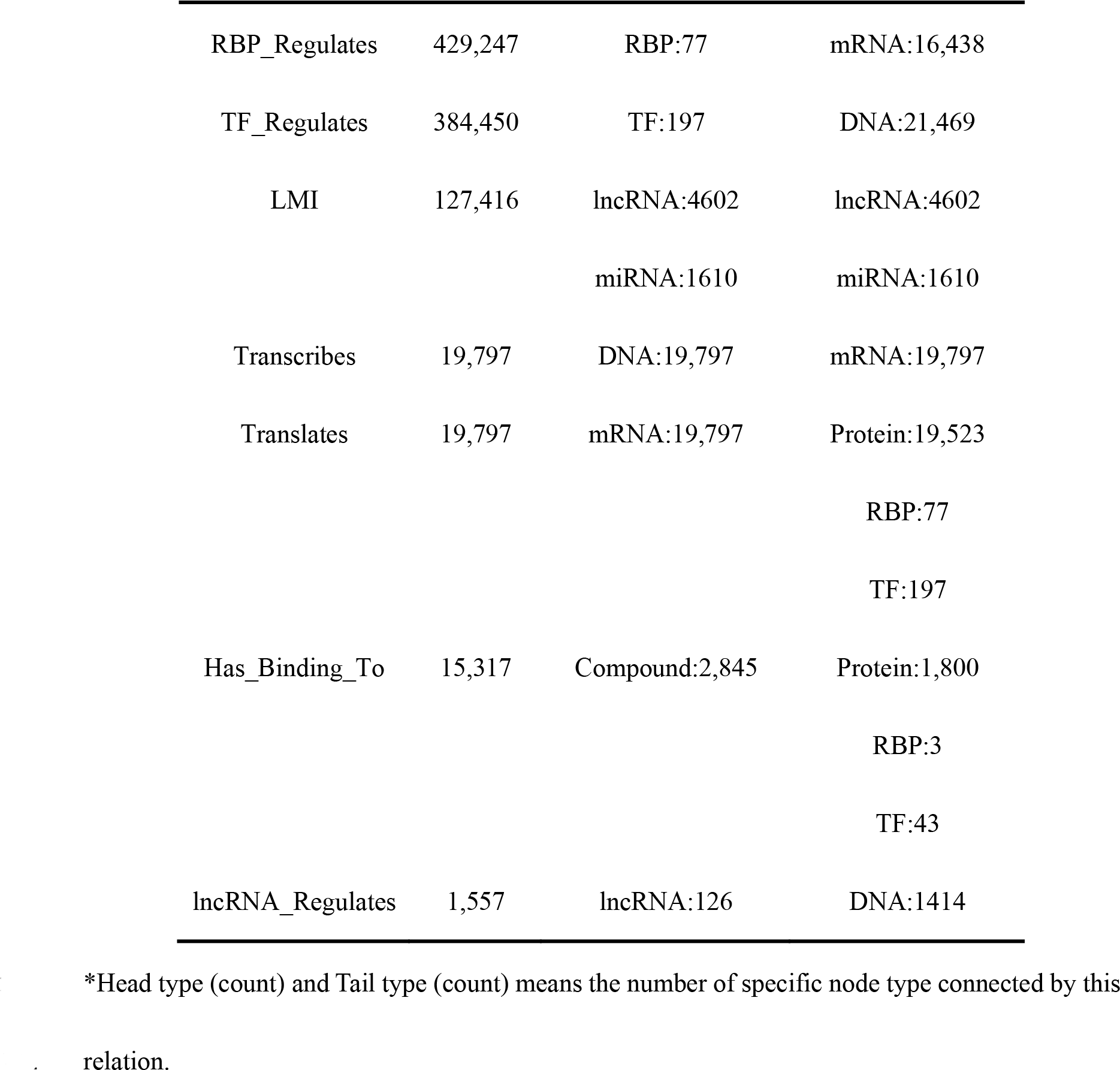
Relations in the chemical perturbation knowledge graph.

| Type of relations | Count | Head type (count) | Tail type (count) |
| --- | --- | --- | --- |
| Upregulates | 2,178,400 | Compound:10,892 | mRNA:10,166 |
| Downregulates | 2,178,400 | Compound:10,892 | mRNA:10,166 |
| miRNA_Regulates | 970,099 | miRNA:618 | mRNA:12,544 |
| PPI | 497,282 | Protein:16,299 | Protein:16,299 |
|  |  | RBP:77 | RBP:77 |
|  |  | TF:194 | TF:194 |
| RBP_Regulates | 429,247 | RBP:77 | mRNA:16,438 |
| TF_Regulates | 384,450 | TF:197 | DNA:21,469 |
| LMI | 127,416 | lncRNA:4602 | lncRNA:4602 |
|  |  | miRNA:1610 | miRNA:1610 |
| Transcribes | 19,797 | DNA:19,797 | mRNA:19,797 |
| Translates | 19,797 | mRNA:19,797 | Protein:19,523 |
|  |  |  | RBP:77 |
|  |  |  | TF:197 |
| Has_Binding_To | 15,317 | Compound:2,845 | Protein:1,800 |
|  |  |  | RBP:3 |
|  |  |  | TF:43 |
| lncRNA_Regulates | 1,557 | lncRNA:126 | DNA:1414 |

\*Head type (count) and Tail type (count) means the number of specific node type connected by this relation.

### Train and inference

As illustrated in Fig. 1C, the training of PertKGE involves several steps. (1) Random initial embedding: entities and relations in the knowledge graph are represented in low-dimension space by embedding using Glorot initialization^37^. (2) Negative sampling: for each existing triple, 100 corrupted triples are randomly generated using the Bernoulli negative sample strategy^38^. (3) Triples scoring: The DistMult is used as a scoring function to assess the validity of existing triples and corrupted triples. (4) Optimization: the margin loss is computed to maximize the scores of existing triples, minimize the scores of corrupted triples, and update embeddings. This process iterates for n times until the embeddings of entities and relationships optimally represent the semantics of chemical perturbation.

During the inference stage, users can query PertKGE with a compound or target of interest, depending on their objective, such as target inference or ligand VS. Following the query, PertKGE calculates the CPI scores using the trained KGE and generates a recommendation list based on these scores. The top N candidates within the recommendation list are typically chosen for further experimental validation. In the subsequent sections, we aim to evaluate the effectiveness of PertKGE and its integration into the drug discovery stage (Fig. 1D).

### PertKGE enables accurate and robust target inference in a compound cold-start setting

This work investigates the effectiveness of PertKGE for target inference in compound cold- start settings, where compounds lack known CPI annotations rather than those without any information^39^. Similarly, “target cold-start” refers to targets without CPI annotations in this context. As shown in Fig. 2A, we focus on two compound cold-start scenarios: (1) In current knowledge graph, 74% of compounds have DEGs from LINCS Phase I but lack CPI annotations. In this case, direct queries within the knowledge graph suffice. (2) In most cases, query compounds are not included in the knowledge graph. This should be addressed by adding the compound-induced DEGs to the knowledge graph before querying.

**Fig. 2.**
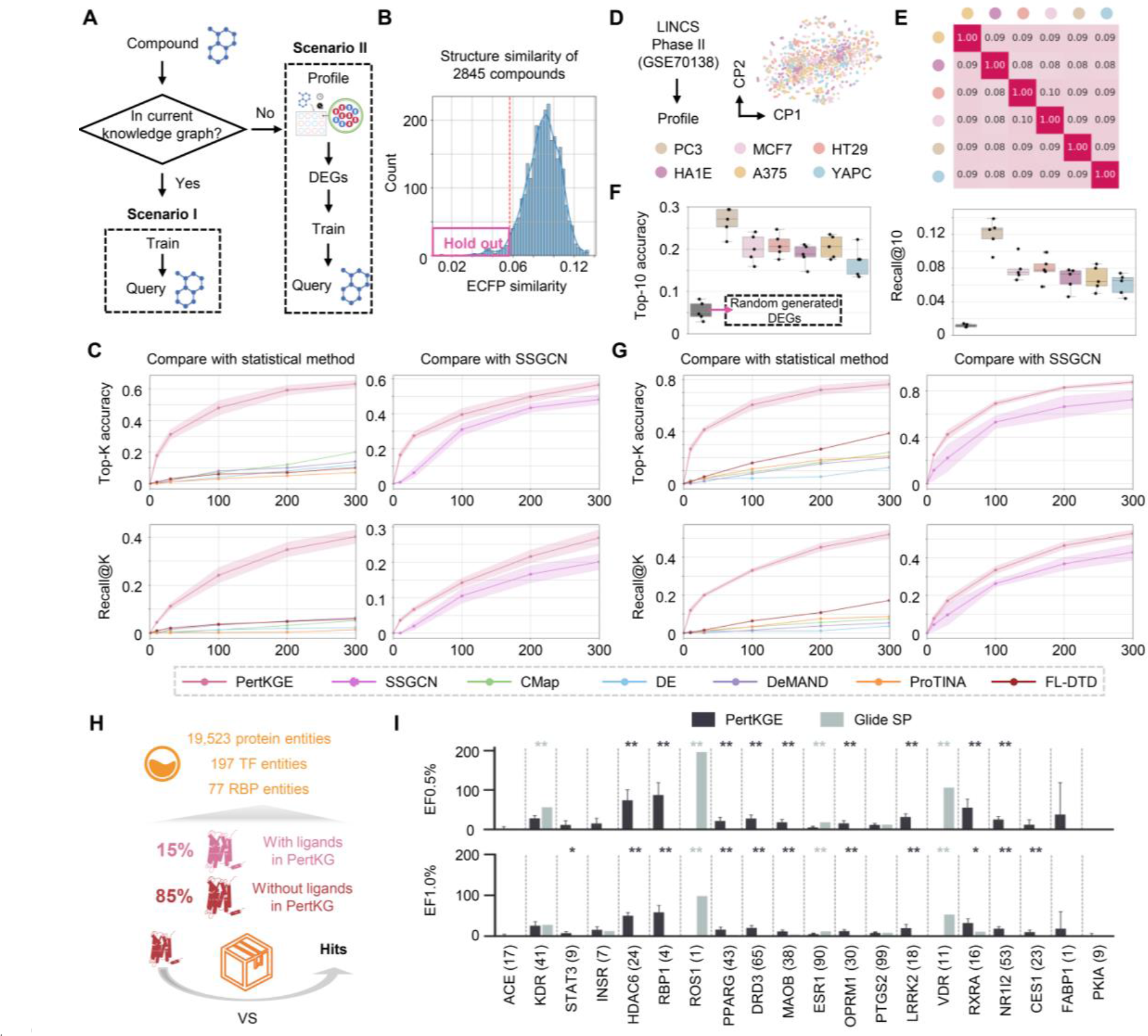
Evaluating CPI prediction performance. A, Illustrations of two compound cold-start scenarios. B, Selection of 100 compounds with the most significant structural differences among all compounds with known targets, by calculating the mean extended-connectivity fingerprints (ECFP) similarity with other molecules. C, Targets inference performance evaluation with PertKGE, SSGCN and other baseline methods in scenario I. The solid line represents the average value of the results from five-fold cross-validation, and the shaded area indicates the range of standard deviation. D, Dimensionality reduction visualization of 170 compounds’ CGS from 6 cell lines in LINCS Phase II. E, Comparison of the 170 compounds’ most upregulated DEGs in different cells by calculating their Tanimoto coefficients. F, Comparison of the target inference performance for 170 compounds’ DEGs in 6 cell lines and randomly generated DEGs. G, Targets inference performance evaluation with PertKGE, SSGCN and other baseline methods in scenario II. H, Illustration of target cold-start scenario. I, Evaluation of ligand virtual screening performance with PertKGE and Glide- SP. The x-axis represents 20 targets for prediction, with the number of ligands collected for each. The statistical significance level was set as *P < 0.05, **P < 0.01.

In the both scenarios, we follow previous work^21^ using Top-K accuracy to evaluate the proportion of tested compounds for which any true target can be correctly predicted among the top K candidates. In addition, considering polypharmacology (A drug acts on multiple targets), we also use Recall@K, a metric commonly used in recommendation systems, to measure the recall among the top K candidates.

For the first scenario, a hold-out strategy was used to create a test set by masking CPI annotations for 100 compounds with the most significant structural differences (Fig. 2B). The remaining knowledge graph was trained using five-fold cross-validation and tested on the hold-out set. We first compare PertKGE to statistical methods reliant on specific prior assumptions, such as CMap, DeMAND, ProTINA, FL-DTD, and differential expression analysis (DE). As shown in Fig. 2C, PertKGE significantly outperformed these methods in both Top-K accuracy and Recall@K metrics. We also compared PertKGE with SSGCN, another deep learning-based methods. It is noteworthy that SSGCN requires gene knockout signatures for target inference, limiting its applicability to 3832 targets. For a fair comparison, only CPI annotations involving these targets were used for PertKGE’s training. The results showed that PertKGE also significantly outperformed SSGCN in two metrics.

For the second scenario, signatures from LINCS Phase II were used to provide additional compound-induced DEGs. The impact of cell line on target inference was investigated. As depicted in Fig. 2D, CGSs for 170 new compounds across 6 cell lines using the same methodology as in LINCS Phase I, and the top 200 upregulated and downregulated DEGs were extracted. The DEGs from different cell lines exhibited minimal similarity, with an average of 32 intersecting genes among the upregulated and 34 among downregulated DEGs (Fig. 2E, Fig. S2). Subsequently, we assessed the DEGs from six cell lines. As expected, PC-3 cells, whose transcriptional data is the basis for the knowledge graph, yielded the highest performance, with a Top-10 accuracy of 0.266 ± 0.029 and a recall@10 of 0.120 ± 0.016. Notably, PertKGE exhibited predictive capabilities across other cell lines despite their dissimilarities (Fig. 2F), and this consistency increased with K (Fig. S3). At K=100, almost all cell lines achieved similar performance to that of PC-3. This suggests that PertKGE may have learned expression patterns that are independent of cell context. To compare, we generated the same number of DEGs for each compound randomly. When these DEGs were used, the model’s performance dropped significantly. However, it still retained some predictive power, potentially due to biases in the datasets (Fig. S3). Based on the results above, we selected the DEGs from PC-3 for comparison with statistical and deep learning methods. Consistent with previous findings, PertKGE also significantly outperformed them (Fig. 2G).

### PertKGE demonstrates promising VS capabilities in the target cold-start setting

VS for targets without any ligands is a significant but challenging scenario in drug discovery. This means ligand-based drug design (LBDD), such as ligand structural similarity-based search, quantitative structure-activity relationship (QSAR), and pharmacophore modelling, cannot be directly applied. As shown in Fig. 2H, 85% of targets, including RBP, TF and Protein entities, in the knowledge graph do not have any known ligands. Hence, it is very valuable to evaluate the ability of PertKGE to screen hits from 10,892 compounds in the knowledge graph for these targets.

Most transcriptomic-based CPI prediction methods are primarily used for target inference. While FL-DTD and SSGCN have reported applications in VS, the former lacks relevant implementation on its provided website, and the latter is unsuitable for target cold-start settings (Fig. S4). Here, we compared PertKGE with the most prevalent structure-based drug design (SBDD) method, molecular docking. Glide-SP^40^, known for its powerful VS capabilities, is selected as a baseline. We also used a hold-out strategy, masking the CPI annotations of 20 targets with 3D- structure in Protein Data Bank (PDB)^41^. Then, we trained PertKGE using five-fold cross-validation strategy and tested on the hold-out set. Note that we selected targets with available 3D structures for ease of comparison with Glide-SP. However, PertKGE can perform VS for targets without 3D structures.

As shown in Fig. 2I, we evaluated the VS capabilities of PertKGE and Glide-SP with 20 targets and 10,892 compounds. Consistent with SBDD work^42^, we use the enrichment factor (EF) as a performance metric. In terms of EF0.5%, PertKGE significantly outperforms Glide-SP in 9 targets, while Glide-SP performs better in 4 target. There is no significant difference between PertKGE and Glide-SP in the remaining 7 targets. Regarding the EF1.0%, PertKGE demonstrated higher virtual screening capabilities, outperforming Glide-SP in 11 targets, while Glide-SP performs better in 3 targets. There is no significant difference between PertKGE and Glide-SP in the remaining 6 targets. For the Reactive Oxygen Species (ROS), Glide-SP showed high enrichment capabilities. However, this is because, out of 10,892 molecules, only one was a ligand of ROS, and Glide ranked it fourth. In summary, PertKGE demonstrated better performance than Glide-SP in most targets when VS against these 10,892 compounds.

### Multi-level regulatory events are essential to alleviate dataset bias

In the previous experiments, we observed a counter-intuitive result that PertKGE still exhibited some predictive capability even when using randomly generated DEGs. Actually, this is a common limitation of ML models, where they tend to assign high scores to entities that are over-represented in the training set, leading to biased predictions^43^. This limitation deviates from our goal of finding a reliable mapping from compound-induced DGEs to CPI. In this work, we introduce multi-level regulatory events to strengthen the connections between genes, alleviating this limitation. To illustrate, we designed a test where the model was only allowed to use compound-induced DEGs for prediction (Method).

Initially, we attempted a comparison with other commonly used BKGs^25,26,28^. However, such a direct comparison is not accurate because many understudied compounds can only be incorporated as isolated nodes in other BKGs, leading to substandard performance. Instead, we pruned other BKGs to replace the process component of our knowledge graph, while keeping the cause and effect components unchanged (Fig. 3A). For convenience, we still refer to them by their original names. Fig. 3B shows the process component derived from different BKGs. It is clear that our knowledge graph represents genes in various forms such as DNAs, mRNAs, TFs, RBP and so on. In contrast, in other BKG, genes are typically represented in only one form, like proteins in BioKG and PrimeKG, or genes in HetioNet. Furthermore, while they have triples of the same order of magnitude, our knowledge graph captures regulatory events between different forms of genes. However, BioKG, HetioNet, and PrimeKG primarily describe relationships between genes and coarse-grained nodes such as biological processes and pathways, as well as nodes less related to chemical perturbation such as diseases and anatomies.

**Fig. 3.**
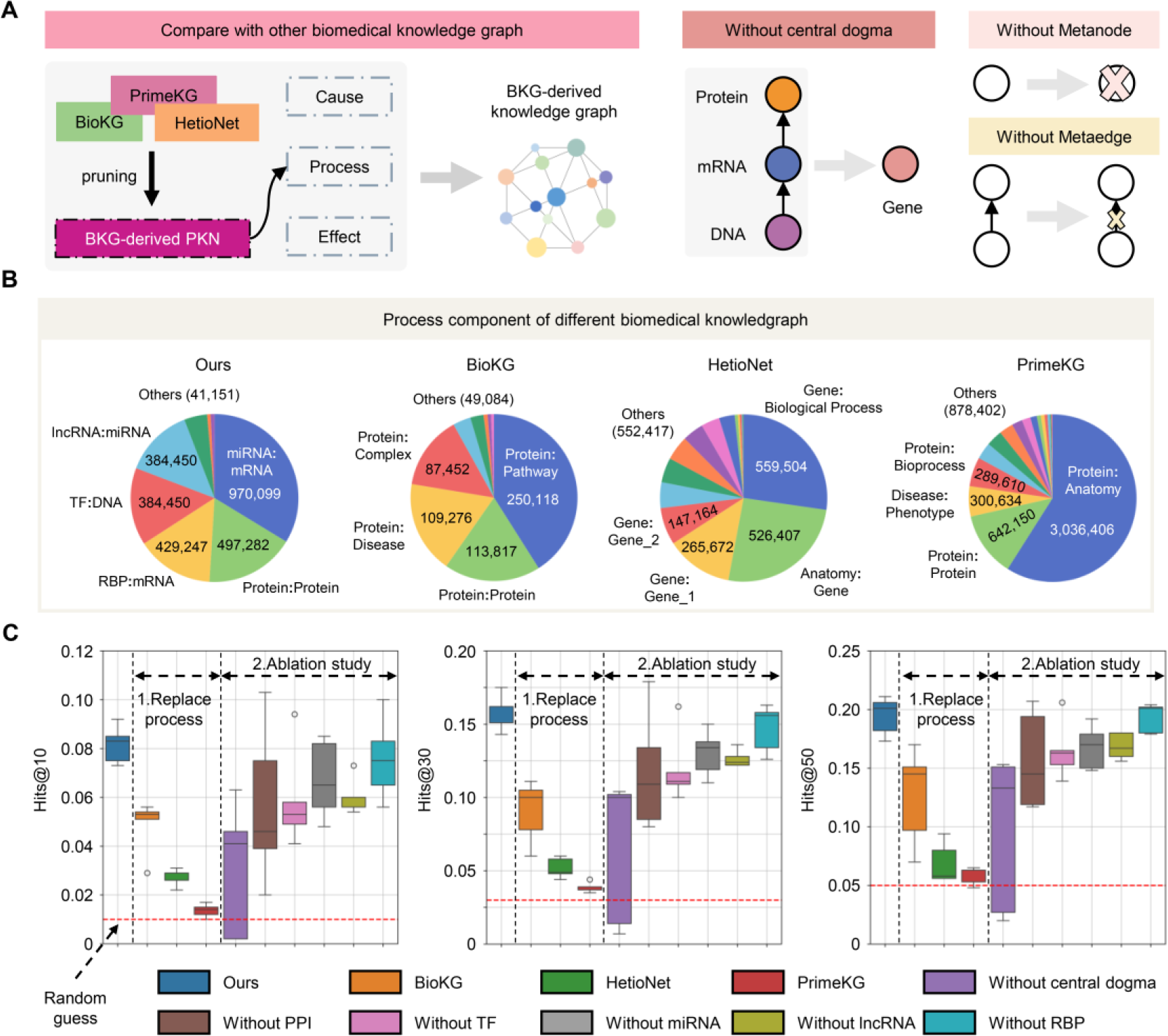
Study of multi-level regulatory events. A, Compare with other BKGs and ablation studies in unbiased test. Left, construction of BKG-derived knowledge graphs. BKGs were Pruned by removing drug entities and drug related triples. Right, illustrations of three kinds of ablation studies: without central dogma, without metanode and without metaedge. B, The pie charts illustrate the relations in different process components. Since different BKGs use different descriptions, we uniformly represent relations in the form of head:tail. If there are multiple different head:tail relationships within the same BKG, we assign them numerical identifiers like head:tail_1. C, Performance of CPI prediction using different process components. Red dashed line means the performance of random guess.

We trained three alternatives using the same approach as PertKGE. As shown in Fig. 3C, PertKGE outperformed the three alternatives across all metrics. This implies that process component based on regulatory events between different forms of genes strengthen the connection between cause-and-effect components by sharing the same context of chemical perturbation. Although other BKGs may also include entities related to chemical perturbation, such as pathways and biological processes, their descriptions are typically coarse-grained (manifested as a pathway or biological process connecting to multiple gene in the graph). This makes it difficult for the model to learn finer regulatory patterns. Interestingly, when we use PrimeKG, despite having the most training triples, it performs nearly as poorly as random guessing. This might because PrimeKG contains over 3 million triples describing relationships between proteins and anatomies, which are largely irrelevant to perturbation. This redundancy knowledge may even impede the model’s ability to learn other useful knowledge, resulting in poor performance.

We further explored which regulatory events most significantly contribute to the performance of PertKGE through an ablation study (Fig. 3C). It can be found that decoupling genes into DNAs, mRNAs, TFs, RBPs, and other proteins, in accordance with their roles in the central dogma, significantly improves the model’s performance. This improvement may be due to the restoration of the biological system’s hierarchical structure, enabling the model to differentiate the semantics of various regulatory events. However, this consideration is often overlooked in other BKGs. Furthermore, in line with previous studies^16,22^, both PPI and TF-mediated regulatory events indeed enhance gene connections. Removing these regulatory events results in a significant performance decrease. In this study, we modeled the impact of other regulatory elements (RBP, miRNA, lncRNA) on gene expression for the first time. The results indicated that integrating regulatory events based on miRNA and lncRNA enriched the model’s understanding of the biological regulatory network, thereby further enhancing the model’s predictive capabilities. However, the addition of regulatory events based on RBP only led to a slight improvement. This could be since RBPs exert a more refined regulatory role in biological networks^44,45^. In the knowledge graph, each RBP is associated with an average of 5000 downstream genes, but only one type of relation is used to describe this regulation.

### Secondary pharmacology study of K-756 by PertKGE

K-756, a Wnt/β-catenin pathway inhibitor targeting tankyrase (TNKS), is currently in preclinical testing. It selectively inhibits the ADP-ribosylation activity of TNKS1 and TNKS2 with IC50 values of 31 nM and 36 nM, respectively^46^. XAV-939, another preclinical TNKS inhibitor, inhibits TNKS1 and TNKS2 with IC50 values of 11 nM and 4 nM, respectively52. In our studies, we accidentally discovered that K-756 exerts unique anti-tumor immune activity in the 4T1 orthotopic breast cancer mouse model, in contrast to XAV-939. This was demonstrated by K-756 significantly increasing the infiltration of CD3+ T cells in tumors, the frequency of CD8+ cytotoxic T cells within CD3+ T cells, and reducing the expression of CD8+ T cell exhaustion marker PD-1 (Fig. 4A, Fig. S5A-C). However, XAV-939 administration did not result in a noticeable change in the infiltration of CD3+, CD8+, and PD-1+CD8+ T cells in tumors (Fig. 4A, Fig. S5A-C). Notably, we also observed that K-756 exhibited an obvious stronger potency to inhibit tumor growth than XAV-939 (Fig. S5D). These results have encouraged us to explore the secondary pharmacology of K-756, to elucidate its mode of action not related to TNKS and explain why it exerts the unique anti-tumor immunotherapy effects.

**Fig. 4.**
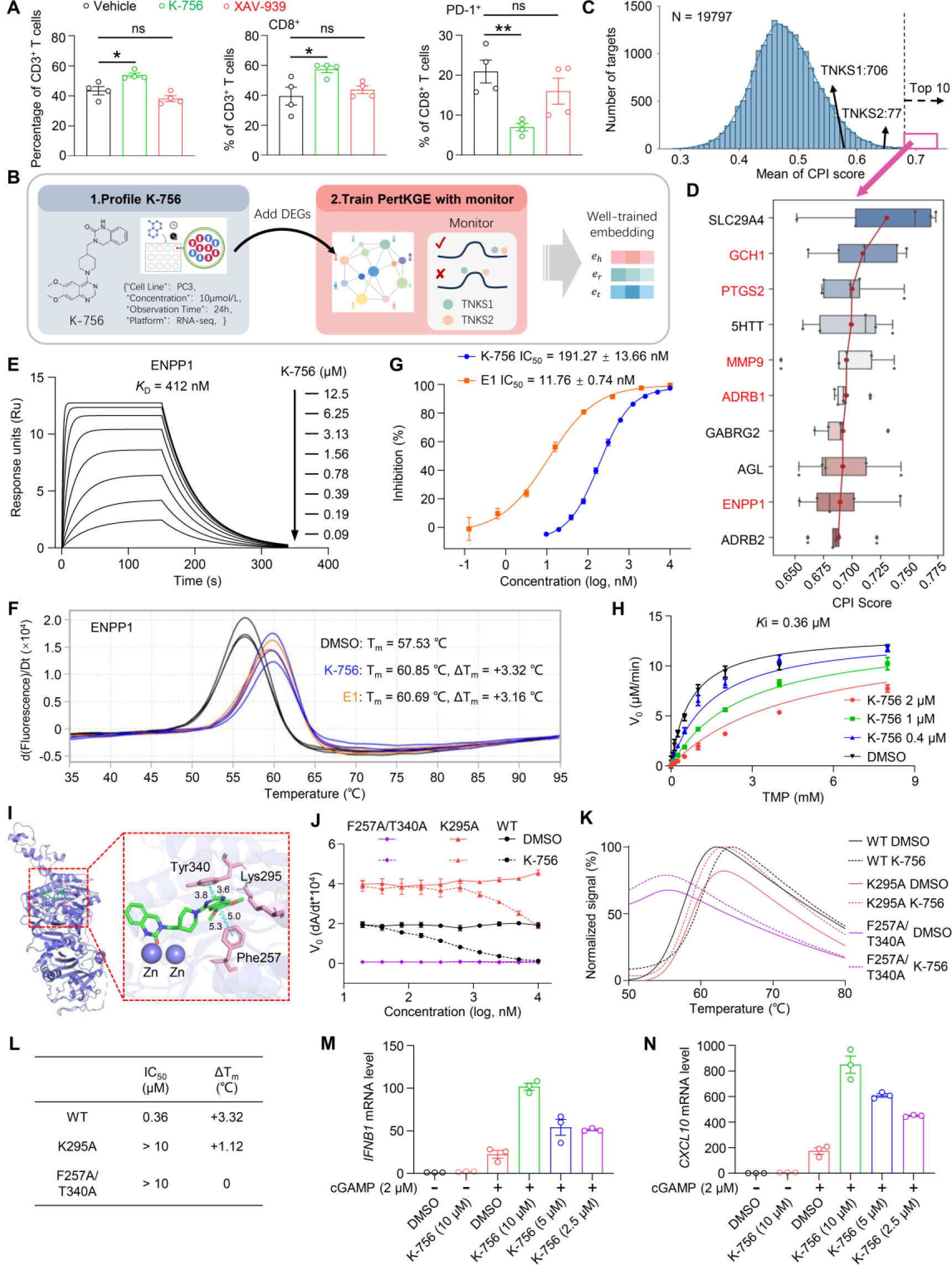
Secondary pharmacology study of K-756 by PertKGE. A, Impact of K-756 and XAV- 939 on the infiltration of CD3+, CD8+, and PD-1+CD8+ T cells in tumors was assessed by flow cytometry (n=4). BALB/c mice were orthotopically inoculated with 4T1 breast cancer cells and administered 30 mg/kg K-756 or XAV-939 daily via intraperitoneal injection. B, Pipeline of target inference for K-756. C, The distribution of predicted scores for all 19797 targets. D, Top 10 predicted targets for K756 are presented in a box plot, illustrating the results of five-fold cross- validation. Rankings are based on the mean predictions from cross-validation. Targets highlighted in red are associated with tumor immunity. E, Binding affinity measurement of K-756 to ENPP1 protein using SPR assay. Graphs depicting equilibrium response units versus K-756 concentrations were plotted. F, Impact of K-756 and E1 on the thermal stability of ENPP1 protein, as determined by protein thermal shift (PTS) assay. G, Dose-dependent inhibition of K-756 and E1 against ENPP1. The substrate for the ENPP1 enzymatic reaction is thymidine 5′-monophosphate p-nitrophenyl ester (TMP). H, The steady-state enzyme kinetics analysis of ENPP1 was conducted in the presence of various concentrations of K-756. I, Docking results for K-756 using a reported ENPP1 X-ray crystal structure (PDB entry 6WEU) as the template. The figures were generated using PyMOL (http://www.pymol.org/). J-L, The impact of K-756 on enzymatic activity and thermostability of mutant ENPP1 protein. M-N, IFNB1 and CXCL10 mRNA levels in THP-1-derived macrophages were measured following treatment with 2 μM cGAMP alone, or 2 μM cGAMP combined with various concentrations of K-756 for 12 h. Error bars indicate the mean ± SEM of three biologically independent experiments (G, H, J, M, N). A two-tailed unpaired t-test was used to analyze significant differences between groups (*, P < 0.05; **, P < 0.01; ns, no statistical difference, P > 0.05).

As K-756 is not present in the knowledge graph, its target inference falls into the second scenario discussed earlier. As illustrated in Fig. 4B, we first measured the transcriptional profile of K-756-treated PC-3 cells (the differential gene analysis results is provided in Fig. S6) and converted DEGs into triplets before adding them to knowledge graph. We then trained PertKGE to test if it could predict the known targets of K-756. As Fig. 4C shows, PertKGE ranked TNKS1 at 706th and TNKS2 at 77th among 19797 candidates, which indicates that the trained embeddings capture the relationships between the DEGs and targets of K-756. Then we focus on the top 10 predicted targets of K-756 (Fig. 4D), which represent the targets that PertKGE identified as the most likely to bind with K-756. GTP cyclohydrolase 1 (GCH1)^47^, prostaglandin-endoperoxide synthase 2 (PTGS2)^48^, matrix metalloproteinase 9 (MMP9)^49^, adrenergic receptor beta 1 (ADRB1)^50^, and ENPP1^51^, which have been reported to be associated with anti-tumor immunotherapy, were selected for the subsequent analyses. We evaluated the binding affinity of these proteins with K-756, excluding ADRB1 as it is not readily available. Notably, ENPP1 demonstrated a nanomolar binding affinity towards K-756, evidenced by a KD constant of 412 nM, measured using a surface plasmon resonance (SPR) assay (Fig. 4E). However, no binding interaction was observed between K-756 and the other three proteins (Fig. S5E-G). ENPP1 knockout or pharmacological inhibition prevents the hydrolysis of tumor-derived cGAMP, leading to the accumulation of cGAMP and the reduction of adenosine in the tumor microenvironment. This activates the STING signaling pathway and relieves adenosine-mediated immune suppression, ultimately exerting anti-tumor immune effects^51,52^. Based on these findings, we purified recombinant ENPP1 protein and further investigated K-756 as a potential inhibitor of ENPP1. K-756 significantly increased the thermal stability of the ENPP1 protein (Fig. 4F), indicating a direct binding interaction with ENPP1. The half-maximal inhibitory concentration (IC50) of K-756 against the enzymatic activity of ENPP1 was 191.27 nM (Fig. 4G). For comparison, ENPP1-IN-1 (E1, WO2019046778), used as a positive control, exhibited an IC50 value of 11.76 nM in inhibiting ENPP1 enzyme activity. Classic steady- state enzyme kinetic experiments showed that as the concentration of substrate TMP increased, Vmax remained constant while Km increased (Fig. 4H), suggesting that K-756 competes with the substrate for binding to ENPP1. Docking-based molecular simulations showed that K-756 inserts into the substrate-binding pocket. The pyrimidine ring of K-756 forms π-π stacking interactions with Phe257 and Tyr340, and hydrogen bonds with Lys295, firmly anchoring K-756 in the pocket (Fig. 4I). Furthermore, 100 ns molecular dynamics simulations revealed a stable conformation between K-756 and ENPP1, with sustained interactions observed between K-756 and Phe257, Lys295, Thr340, and Tyr371 (Fig. S5H and S5I). To confirm this binding mechanism, we generated two ENPP1 protein mutants, K295A and F257A/T340A. These mutations significantly reduced or completely abolished the binding and the enzymatic inhibitory effect of K-756 on ENPP1 (Fig. 4J- L). Collectively, these results strongly suggest that K-756 binds to the substrate-binding pocket of ENPP1. Furthermore, as expected, K-756 significantly enhanced the transcription of downstream cytokines in the STING pathway, including IFNB1, CXCL10, and IL6, when induced by cGAMP in THP-1-derived macrophages (Fig. 4M and 4N, Fig. S5J).

To determine whether the inhibition of ENPP1 by K-756 is a common characteristic of TNKS inhibitors or a unique feature of the K-756 molecule, we simultaneously tested the inhibitory activity of K-756 and three other TNKS inhibitors: VAX-939, NVP-TNKS656, and RK-287107, on ENPP1 enzyme activity. As shown in Fig. S5K, only K-756 exhibited inhibitory activity against ENPP1. The unique pharmacological activity of K-756 indicates that dual-target inhibitors of TNKS and ENPP1 may have promising synergistic anti-tumor activity. In summary, the success of repurposing TNKS inhibitor K-756 to ENPP1 inhibitor demonstrated that the practical and promising targets inference ability of PertKGE.

### PertKGE identified five new scaffold hits for ALDH1B1

Aldehyde dehydrogenases (ALDHs) are highly expressed in multiple cancer types, contributing to cancer progression, therapy resistance, and immune evasion^53^. ALDH1B1, a mitochondrial ALDH isoform, promotes colorectal and pancreatic cancer^54^. Selective pharmacological inhibition of ALDH1B1 has been shown to hinder the growth of colon cancer spheroids and patient-derived organoids^54,55^. Moreover, the viability of Aldh1b1-knockout mice suggests that blocking ALDH1B1 has tolerable effects on normal physiology^54,56^. These results indicate that ALDH1B1 is a promising cancer drug target. To our knowledge, imidazoliums and guanidines^54^ are the only effective ALDH1B1 inhibitors. However, they lack drug-like properties and are primarily used as molecular probes to study ALDH1B1 functions. Therefore, there is an urgent need to discover novel scaffold ALDH1B1 inhibitors for cancer treatment.

Here, PertKGE was utilized as a virtual screening tool to identify novel ALDH1B1 inhibitors. As Fig. 5A shows, screening was conducted on 7,403 small molecules in the knowledge graph after filtering out those may be pan-assay interference compounds (PAINS) and with heavy molecular weight, and the top 100 candidates predicted by PertKGE were purchased from commercial libraries for further validation.

**Fig. 5.**
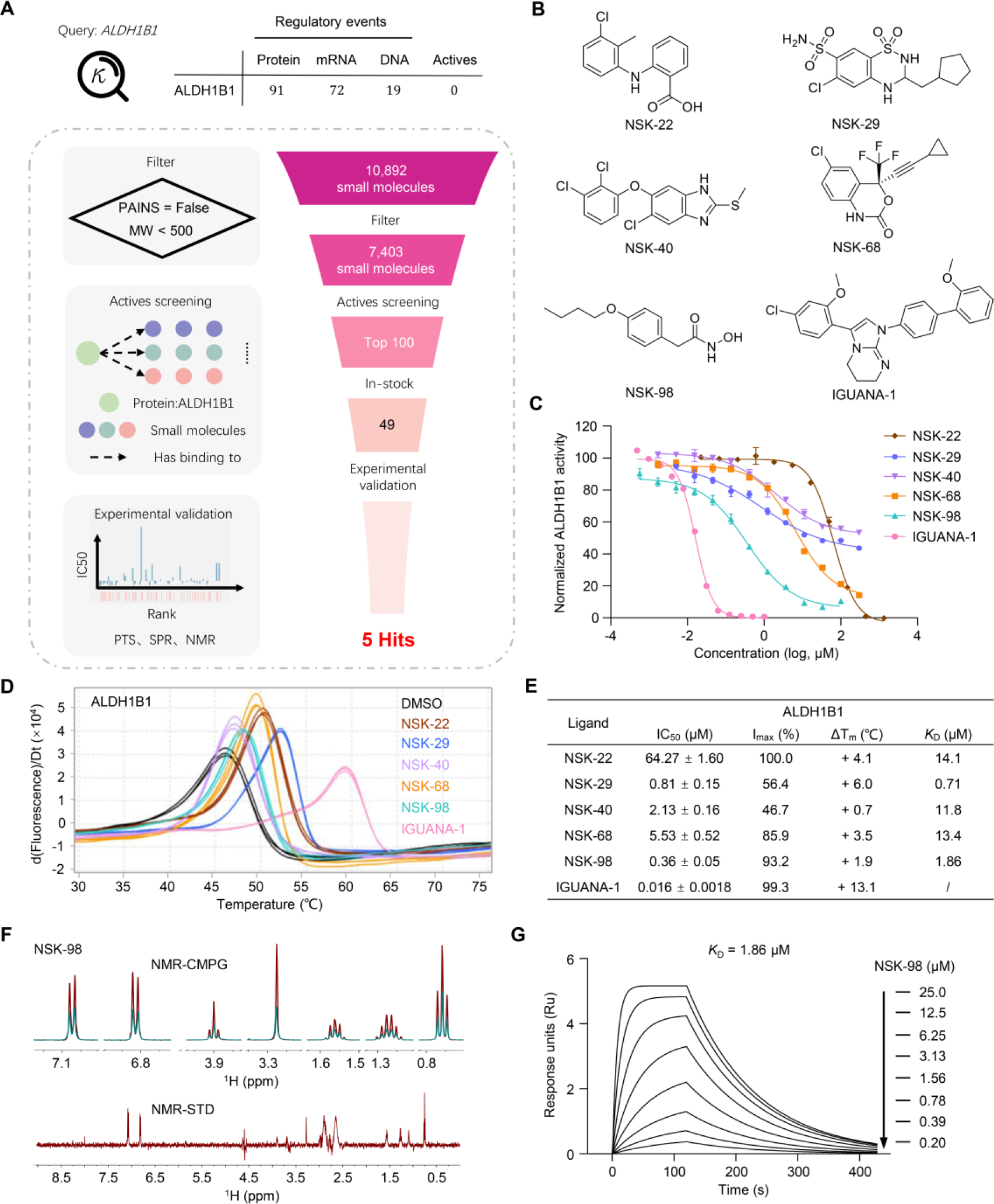
PertKGE identified five new scaffold hits for ALDH1B1. A, The query for ALDH1B1 in the current PertKGE reveals three existing forms of ALDH1B1, and there is no known active molecule targeting protein-level ALDH1B1. The dashed box outlines the scheme of the active screening protocol for ALDH1B1. B, Chemical structures of five hit compounds: NSK-22, NSK- 29, NSK-40, NSK-68, NSK-98, and reported ALDH1B1 inhibitor IGUANA-1. C, Dose–response curves of NSK-22, NSK-29, NSK-40, NSK-68, NSK-98, and IGUANA-1, determined by ALDH1B1 enzyme kinetics assay. Error bars represent the mean ± SEM of three independent experiments. D, Impact of NSK-22, NSK-29, NSK-40, NSK-68, NSK-98, and IGUANA-1 on the thermal stability of ALDH1B1 protein, as determined by PTS assay. E, Summary of IC50 values of the indicated compounds determined by enzyme kinetics assays, melting temperature differences (ΔTm) determined with PTS assay, and KD values measured via SPR assay. F, Nuclear magnetic resonance (NMR) measurement of direct binding between NSK-98 and ALDH1B1 protein. CPMG NMR spectra for NSK-98 (red), NSK-98 in the presence of 5 μM ALDH1B1 protein (green). The STD spectrum for NSK-98 was recorded in the presence of 5 μM ALDH1B1 protein. G, Binding affinity measurement of NSK-98 to ALDH1B1 protein using SPR assay. Graphs depicting equilibrium response units versus NSK-98 concentrations were plotted.

Initially, an enzyme kinetics assay was carried out to measure the inhibitory activity of the predicted actives against ALDH1B1, with IGUANA-1^54^ as a positive control due to its potent inhibitory activity and commercial availability. Out of the 49 commercially available candidates, NSK-22, NSK-29, NSK-40, NSK-68, and NSK-98, showed significant inhibitory activity on ALDH1B1, at a hit rate of 10.2% (Fig. 5B, C, and E). Further, the impact of these five compounds on the thermostability of recombinant ALDH1B1 protein was evaluated. All compounds substantially improved the thermal stability of ALDH1B1 protein (Fig. 5D and E), suggesting these compounds could directly bind to ALDH1B1. Additionally, the attenuation of NSK-22, NSK-29, NSK-68, and NSK-98 signals after incubation with ALDH1B1 protein was observed in the Carr-Purcell-Meiboom-Gill (CPMG) nuclear magnetic resonance (NMR) spectra. Positive saturation transfer difference (STD) signals in the STD spectrum were also noted (Fig. 5F, Fig. S7A-C), further indicating their direct binding to ALDH1B1. NSK-40 was not included in this NMR analysis due to its poor solubility in the assay buffer, preventing signal collection. To determine the binding affinity between these five hits and ALDH1B1, a SPR assay was conducted. The results showed that the binding affinity of the five hits ranged from 0.71 to 14.1 μM (Fig. 5E and 5G, Fig. S7D-G). Collectively, these results strongly demonstrate that NSK-22, NSK-29, NSK-40, NSK-68 and NSK- 98 can directly bind to ALDH1B1 and inhibit its enzymatic activity. Compared to the current two classes of ALDH1B1 inhibitors, these 5 hits possess novel scaffolds and have a clinical drug history (Fig. S8), holding promise for further development.

We also examined whether these hits could be identified with conventional SBDD approach. However, molecular docking ranked them at 1937th, 2145th, 1509th, 6488th, and 7322th, respectively. (Fig. S7H). This showed that PertKGE could be an excellent virtual screening, discovering actives overlooked by conventional methods.

## Discussion

Exposure of cells to small molecules often triggers multi-level remodeling, which can be observed through perturbation omics data. These data provide a dynamic and more realistic view of the impact of compounds on cells, making it a promising information source for understanding CPI. In this study, we developed a novel method, PertKGE, based on a biologically meaningful knowledge graph to systematically mine perturbation transcriptomics. By integrating a range of regulatory events mediated by factors such as TFs, RBPs, other proteins, miRNAs and lncRNAs, and using KGE algorithm, PertKGE can better understand the context of chemical perturbation, leading to accurate and robust CPI predictions. Our method has outperformed baseline methods in two cold-start settings and has avoided pitfall associated with ML. These encouraging computational results led us to incorporate PertKGE into the early stages of drug discovery. We applied PertKGE in two real-word application scenarios: (1) How to explain K-756’s unique anti-tumor immune activity in phenotype-based screening. The experimental validation of the top ten targets recommended by PertKGE strongly suggests that K-756 binds to the substrate-binding pocket of ENPP1, indicating that the development of dual-target inhibitors of TNKS and ENPP1 may serve as a promising synergistic anti-tumor strategy. (2) How to find novel scaffold inhibitors with drug-like properties for a less studied target, like ALDH1B1. Through the combined use of PertKGE and experimental methods, we identified 5 hits with a 10.2% hit rate. These hits possess novel scaffolds, indicating potential for further development. The success of PertKGE both in computational experiments and practical applications demonstrates its potential as a promising tool for helping pharmacologists in understanding the MOA of compounds and screening promising inhibitors.

In drug discovery, the use of a knowledge graph to integrate and analysis multi-omics data is a promising approach. Unlike previous methods that used compound-induced transcriptomic profiles as the primary representation of nodes^27^ or simply added up/down-regulation relations in existing BKGs^25^, our work underlines causal reasoning. We introduce a new knowledge graph that decomposes chemical perturbations into three components: a cause component made up of CPI, a process component consisting of multiple regulatory events, and an effect component comprising observed DEGs. This approach customizes the entire knowledge graph for a specific context, namely chemical perturbation, and has shown effectiveness in both computational analyses and experimental validations. We believe that CPI prediction is a significant challenge, and we are far from a complete solution. In the future, the knowledge graph we created could be further enhanced by considering more and finer regulatory events. Additionally, the application scope can be broadened by using PertKGE to analyze other types of large-scale omics data, such as perturbation proteomics^57^ and cell images^58,59^.

## Methods

### Constructing the chemical perturbation knowledge graph with data from multiple database

We downloaded relevant raw data from multiple domain databases, subsequently converting them into a standard triple format denoted by (*head*, *relation*, *tail*). To establish connections between triples originating from distinct database sources, we employed entity alignment. Finally, the Networkx python package^60^ was used to retain the largest connected subgraph through pruning.

**PubChem**^61^. Managed by the National Center for Biotechnology Information (NCBI), PubChem serves as a comprehensive repository of chemical information. The diverse representation of each chemical substance is standardized through the PubChem CID (Compound Identifier), offering a consistent reference for a specific substance across different contexts and datasets. In our study, we employed PubChem’s online service to convert all compounds into their corresponding CIDs using Simplified Molecular Input Line Entry System (SMILES).

**UniProt**^62^. UniProt is a comprehensive, freely accessible database providing detailed information on protein sequences and functional annotation. Our acquisition of gene names for human proteins was conducted through the official UniProt website. Subsequently, we established a correlation between the acquired gene names and primary names. In chemical perturbation knowledge graph, all proteins are uniquely identified and represented by their primary names.

**STRING**^33^. STRING serves as a repository consolidating PPIs derived from both experimentally confirmed discoveries and anticipated outcomes. The human PPI network utilized in our study was extracted from the STRING v11.5 database. We converted PPI interactions with a combined score surpassing 700 into bidirectional relationships. Subsequently, we formatted these bidirectional relationships into standard triples for further analysis.

**ENCORI**^29^. ENCORI, previously known as StarBase, is a platform designed for exploring the RNA related interaction networks from CLIP-Seq datasets. We downloaded the interactions mediated by RBP and miRNA-regulated mRNA interactions through the web API and processed them into the standard triple format. Here, hg38 is used as the reference genome, and the chosen interactions must have at least five CLIP-Seq experimental records in the database to ensure accuracy. For miRNA, the alignment was executed based on the gene symbol present in the database.

**RAID**^36^. RAID, a database centered on RNA interactions, is now at its 4.0 version, boasting over 41 million RNA-related interactions spanning 154 different species. Data retrieval from the official website encompassed information on the regulatory effects of lncRNA on DNA, and interactions between lncRNA and miRNA. Our focus was directed towards interactions specific to humans, and we ensured the inclusion of interactions supported by substantial experimental evidence. Subsequently, this data was organized into a standard triple format, aligning lncRNA and miRNA using the gene symbols sourced from the database.

**CHEA**^34^. CHEA, which offers target genes of transcription factors derived from published ChIP-chip, ChIP-seq, and other transcription factor binding site profiling studies, was accessed through the Harmonizome 3.0. We extracted the CHEA dataset and transformed it into a standard triple format, resulting in the acquisition of 384,450 DNA regulatory relationships mediated by 197 transcription factors.

**LINCS**^6^. LINCS, supported by the NIH, systematically captures and documents gene expression patterns across diverse cell lines when subjected to different perturbations under varied experimental settings. The LINCS phase I L1000 dataset (GSE92742, spanning 2012–2015) and the LINCS phase II L1000 dataset (GSE70138, from 2015–2017) were obtained from the Gene Expression Omnibus (GEO) through the Broad Institute. Initially, we opted for the level 5 data, viewing it as a refined depiction of the transcriptional outcomes of a given experiment condition. Focusing on the PC3 cell line’s perturbation data due to its extensive volume, we applied the moderated z-score (MODZ) method to create consensus signatures, capturing essential gene expression changes across different time points and concentrations. Utilizing weights from the original study^6^, we expanded this to cover 10,174 confidently inferred genes. We curated a list of the top 200 genes, comprising both upregulated and downregulated genes, based on corrected expression levels, forming the basis of our triples.

**Other datasets.** In addition to the insights from the databases previously mentioned, we sourced annotation details from LINCS^6^, DepMap^30^, ChEMBL^31^, and DrugBank^32^ to shape cause component of chemical perturbation knowledge graph. Additionally, the central dogma served as prior knowledge for linking DNA to mRNA, and mRNA to proteins.

### Training protocol

The training of PertKG is to embed the entire chemical perturbation knowledge graph into a vector space, thereby obtaining embeddings rich in chemical perturbation semantic for compounds and targets. We denote head entity, tail entity and relation in chemical perturbation knowledge graph as *h*, *t* ∈ *E* (the set of entities) and *r* ∈ *R* (the set of relations). Then PertKG computes validity for each triple (*h*, *r*, *t*) using DistMult as scoring function:

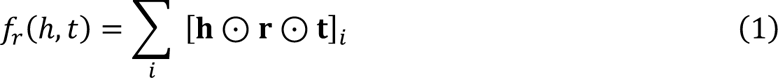

where **h**, **r**, **t** ∈ ℝ*^d^* represent d-dimensional vectors of head entity, relation and tail entity, respectively. Note that for a given entity, its embedding vector is the same when the entity appears as the head or as the tail of a triple. ⊙ denotes element-wise product, and the *i* index is along the feature dimension of the vector resulting from the element-wise production of the different features.

Given training set *S* of existing triples, we minimize a margin-based ranking loss to capture semantic of chemical perturbation knowledge graph:

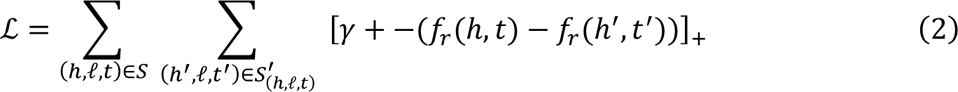

where [*x*]_+_ denotes positive part of *x*, *γ* > 0 is a margin hyperparameter, and

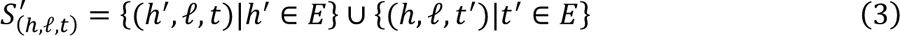

The set of corrupted triples, constructed according to Equation (3), is composed of training triples with either the head or tail replaced by a random entity. To reduce false corrupted triples,

Bernoulli distribution is used to decide to corrupt head or tail^38^.

In this study, five-fold cross-validation strategy is employed in training, wherein the CPI annotations in chemical perturbation knowledge graph have been further partitioned into five folds based on compounds. Four folds are reserved for training PertKG, and the remaining one is used for validation. The whole optimization is carried out by using Adam^63^. Early stopping is used to terminate the training process if the performance of the model on the valid set shows no further improvement. After training, we obtain five well-trained models, and report the mean and standard deviation of the results in the test. The entire training script of PertKG is implement using the TorchKGE^64^.

Additionally, SSGCN was re-implemented adhering to the methodologies outlined in the original paper. The resulting recommendation list was generated based on CPI score of 3832 targets. DeMAND and ProTINA were implemented based on their respective R scripts. ProTINA yielded a descending order of scores for 10,174 target proteins, while DeMAND provided an ascending order of p-values. CMAP and FL-DTD were evaluated using their designated websites. Differential Expression (DE) was executed by sorting the absolute values of gene expression in descending order.

### Evaluation protocol

We evaluated the model’s performance in target inference, ligand virtual screening, and unbiased testing.

**Target inference**: In target inference, we evaluated the performance of identifying targets within the recommendation lists provided by different models. We employed TOP-K accuracy and Recall@K as the metrics.

In the TOP-K accuracy formula, *N* denotes the total number of compounds in test set, *K* is a constant, and *f(i,K)* represents the evaluation of the *i^th^* compound. Specifically, if the compound has at least one target ranked at or below K, the value of *f(i,K)* is set to 1; otherwise, it is set to 0.

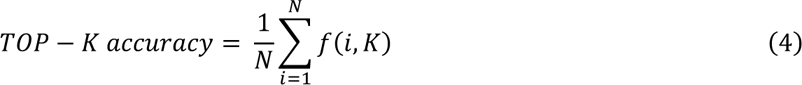

In the Recall@K formula, *Ti* is the number of targets for the *i^th^* compound, and *Ri,k* represents the number of targets ranked at or below *K* for the *i^th^* compound.

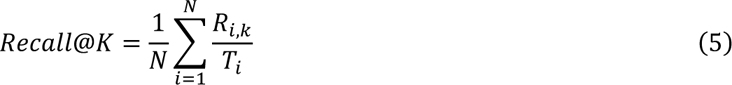

**Ligand virtual screening**: In target inference, we evaluated the performance of identifying ligands within the recommendation lists provided by different models. Our evaluation utilized the Enrichment Factor (EF) as the metric.

In the EF formula, *Hitstotal* is the number of ligands for the target, *Ntotal* is the number of all compounds, *Hitssampled* is the number of compounds among the top N ranked compounds that are active, and *Nsampled* is the number of compounds ranked in the top N.

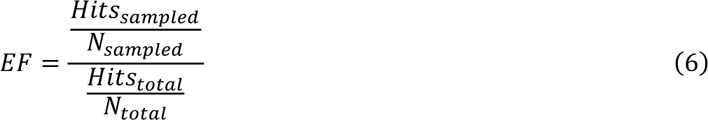

**Unbiased test**: This test was used to evaluate whether the model learned the mapping from DEGs to CPI. In this evaluation, we designated the CPI annotations of 100 compounds for testing purposes. For each reserved CPI, we conducted random sampling to generate 1000 negative compounds. Subsequently, for each positive CPI and its corresponding 1000 negative samples, we assessed the model’s ability to rank the positive compound higher than the negatives. This testing methodology ensures that the model’s performance is contingent on its capacity to learn the mapping from DEGs to CPI. Hits@K was used as metrics in this test.

In the Hits@K formula, *NCPIs* represents the number of compound-protein interaction pairs, and *g(i,K)* represents the evaluation of the *i^th^* CPI pair. If its rank is at or below *K*, the value of *g(i,K)* is set to 1; otherwise, it is set to 0.

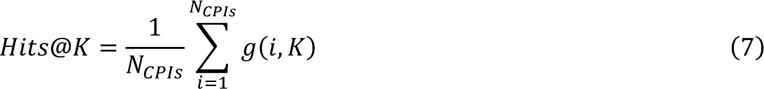

### Recombinant protein expression and purification

For plasmid construction, human ALDH1B1 (residues 20-517) was subcloned into the pET- 15b vector with an N-terminal 6×His tag. To obtain soluble ENPP1 proteins, extracellular region (residues 110-926) of human ENPP1 was fused to the N-terminal secretion signal sequence (residues 1-59) from mouse ENPP2 and cloned into the pcDNA3.1 vector with a Flag tag added to the C-terminal. For expression of ALDH1B1 protein, His-tagged ALDH1B1 plasmid was transformed into BL21(DE3)-Chaperone competent cells (WeiDibio, EM1002S). These cells were subsequently grown in lysogeny broth (LB) medium containing 100 μg/ml ampicillin, 35 μg/ml chloramphenicol and 1 × chaperone inducer at 37°C. Once the optical density at 600 nm reached 0.6∼0.8, the culture was transferred to 16°C, and 0.5 mM isopropyl β-d1-thiogalactopyranoside (IPTG) was added to induce protein expression for 20∼22 hours. The bacteria were then collected by centrifugation (3,000 rpm, 30 minutes, 4°C). Cells were lysed in lysis buffer (Beyotime, P0013Q) containing protease inhibitor cocktail (Beyotime, P1031) and nuclease (Beyotime, D7121-25KU), and the suspension was mechanically rotated at room temperature for 25 minutes. The lysate was centrifuged (16,000 rpm, 30 minutes, 4°C), and the soluble fraction was filtered by 0.22 μm syringe filters and purified on HisTrap columns (Cytiva) using elution buffer (20 mM HEPES, pH 7.4, 500 mM NaCl, 1 mM tris (2-Carboxyethyl) phosphine (TCEP), 500 mM imidazole, and 5% glycerol). The eluted components were exchanged into storage buffer (20 mM HEPES, pH 7.4, 150 mM NaCl) using desalting columns (Cytiva). The purified ALDH1B1 protein was stored in the storage buffer at −80°C. ENPP1 recombinant protein was expressed in suspension Expi293F GnTi-/-cells (Thermofisher, A39240). After 5 days of transfection with the Flag-tagged ENPP1 plasmid, the medium supernatant was collected by centrifuged (16,000 rpm, 1 hour, 4°C) and slowly loaded onto a manually packed Anti-Flag affinity resin. The protein was eluted with an elution buffer consisting of Flag peptides. The elution was further purified on a Superdex 200 increase 10/300 GL column (Cytiva) equilibrated with a buffer containing 20 mM HEPES, pH 7.4, and 150 mM NaCl. The purified ENPP1 proteins were identified using SDS-PAGE and stored at −80°C. For the two ENPP1 protein mutants, K295A and F257A/T340A, the expression and purification methods were identical to those used for the Wild-type ENPP1 protein.

### Enzymatic activity assays

For ALDH1B1, the enzymatic activity assay was conducted in a 384-well white Optiplate (PerkinElmer, 6007290) at room temperature under the following conditions: each well contained 5% (v/v) DMSO, 100 nM His-tagged ALDH1B1 protein, 1 mM nicotinamide adenine dinucleotide (NAD^+^), and 1 mM acetaldehyde in the assay buffer (100 mM sodium phosphate, pH 8.0, 1 mM MgCl2, and 1 mM TCEP). For IC50 determination, His-tagged ALDH1B1 was pre-incubated with NAD^+^ and serially diluted compounds for 5 minutes. Acetaldehyde was then added to each well, and the resulting enzymatic activity was measured based on nicotinamide adenine dinucleotide hydrate (NADH) fluorescence. The fluorescence signal was measured using a TECAN Spark multifunctional microplate reader over the course of 15 minutes with an excitation wavelength of 340 nm and emission wavelengths of 460 nm. For ENPP1, the enzymatic activity assay was conducted in a transparent 384-well plate (NEST, 761001), with a total volume of 50 μL. Each well containing a final concentration of 3 nM Flag-tagged ENPP1 protein, various concentrations of compounds, and 100 μM p-Nph-5’-TMP in the assay buffer (50 mM Tris HCl, pH 8.5, 130 mM NaCl, 1 mM CaCl2, and 5 mM MgCl2). For IC50 determination, Flag-tagged ENPP1 protein was incubated with serially diluted compounds for 10 minutes at room temperature. Subsequently, p- Nph-5’-TMP was added and the absorbance change at 405 nm, indicating the release of p-nitrophenolate, was measured using a TECAN Spark multifunctional microplate reader.

### Protein thermal shift assay

The protein thermal shift assay was conducted using the QuantStudio™ 5 (Applied Biosystems) to evaluate the compound-induced changes in protein thermal stability. For ALDH1B1, His-tagged ALDH1B1 protein (6 μM) was incubated with compounds (100 μM), NAD^+^ (100 μM), and 5 ×SYPRO Orange dye (Sigma, 67-68-5). For ENPP1, Flag-tagged ENPP1 protein (2 μM) was incubated with compounds (50 μM) and 5 × SYPRO Orange dye. The mixtures were then transferred into 384-well plates (Monad, Q50701S) with a final volume of 10 μL. The fluorescence signal was recorded as the temperature was gradually raised from 25°C to 95°C. The data were analyzed using the Protein Thermal Shift™ software v1.4 to determine the Tm value.

### Nuclear magnetic resonance assay

Nuclear magnetic resonance (NMR) assay was performed using a 600 MHz spectrometer (AVANCE III, Bruker) to validate protein-compound interactions. In Carr-Purcell-Meiboom-Gill (CPMG) and saturation transfer difference (STD) NMR experiments, compounds were dissolved to a final concentration of 200 μM in a solution of PBS prepared with D2O, containing 5 μM His-tagged ALDH1B1 protein, 5% DMSO-*d*_6_, and 100 μM NAD^+^.

### Surface plasmon resonance assay

The surface plasmon resonance (SPR) experiments were performed using a Biacore 1K or Biacore 8K instrument (Cytiva) at 25°C. His-tagged ALDH1B1 protein, Flag-tagged ENPP1 protein, GCH1 protein (Cusabio, CSB-EP009317HU), and MMP9 protein (Sino Biological, 10327-H08H) were covalently immobilized onto a CM5 sensor chip (Cytiva) by a standard amine-coupling procedure in 10 mM sodium acetate of different pH (pH 5.0 for ALDH1B1 and MMP9, pH 4.5 for ENPP1 and GCH1). PTGS2 protein (Sino Biological, 12036-H08B) was chelated to the CM5 sensor chip by His capture kit (28995056). The running buffer for ALDH1B1 protein contained 10 mM HEPES, pH 7.4, 400 mM NaCl, 300 μM NAD^+^. The running buffer for ENPP1 and GCH1 protein contained 10 mM HEPES, pH 7.4, 150 mM NaCl. The running buffer for PTGS2 protein contained 10 mM HEPES, pH 7.4, 150 mM NaCl, 0.005% surfactant P20. Compounds were serially diluted into the running buffer and injected onto the sensor chip at a flow rate of 30 μL/minute for 120 or 150 seconds (contact phase), followed by 300 or 180 seconds of buffer flow (dissociation phase). The equilibrium dissociation constant (*K*D) value was derived using Biacore Insight Evaluation software (Cytiva).

### Animal experiment

All animal experiments were conducted with the approval and supervision of the Institutional Animal Care and Use Committee (IACUC) at the Shanghai Institute of Materia Medica, Chinese Academy of Sciences. For the pharmacodynamics experiment, female BALB/c mice (6-8 weeks old) were inoculated with 1×10^6^ 4T1 breast cancer cells into the mammary fat pad. The tumor-bearing mice were randomly divided into three groups when the tumor volume reached approximately 100 mm^3^. Subsequently, the mice were intraperitoneally injected with 30 mg/kg of K-756 (MedChemExpress, HY-U00422) or XAV-939 (MedChemExpress, HY-15147) in a solution containing dimethyl sulfoxide (DMSO) and 20% SBE-β-CD (MedChemExpress, HY-17031) in saline (10/90, v/v) daily for one week. Body weights and tumor volumes of the mice were measured daily. Tumor volumes were calculated using the formula: V= (length×width^2^)/2. At the completion of the study, the mice were euthanized, and the tumors were harvested for further study.

### Flow cytometry analysis

To harvest a single cell suspension, the tumor tissues were shredded with a scissors and treated with digestion solution for 1 hour at 37°C under shaking. The digestion solution was prepared by adding 0.001% hyaluronidase, 0.1% collagenase, 0.002% DNase, 120 μM MgCl2, and 120 μM CaCl2 to RPMI-1640 medium. The digested tumor tissues were filtered through a nylon membrane to obtain a single-cell suspension and treated with ammonium chloride to remove red blood cells. Subsequently, the cell samples were stained with Fixable Viability Stain 700 (BD Horizan, 564997) and the following antibodies: anti-mouse CD16/32 antibody (Biolegend, 101320), CD3-FITC (Invitrogen, 2103752), CD8-BV421 (Biolegend, 100738), PD-1-SB600 (Invitrogen, 2314455). The stained cells were analyzed using the Agilent NovoCyte 3000 instrument. All data were analyzed using the FlowJo software.

### Molecular docking

The molecular docking calculations were based on crystallographic data for the protein structures of ALDH1B1 (PDB:7RAD and 7MJC), and ENPP1 (PDB:6WEU), optimized by the Protein Preparation Wizard at pH 7.0. Subsequently, prepared ligands were docked to the optimized protein using Glide with Standard Precision (SP) mode. All other parameters for the above process were set to default. All docking studies were performed using Maestro of Schrödinger Suites (version 2020-4), and obtained poses were analyzed with PyMOL.

### Molecular dynamics analysis

The molecular dynamics study was performed to examine the conformational changes in the protein that occurred due to the ligand-binding site and to evaluate the effect of these changes over the protein-ligand complex. To evaluate the stability and interaction of the ENPP1 with ligand, simulation study was performed using Desmond Schrödinger Suites (version 2020-4) at 100 ns time period. Water molecules were added to the docking complex of the ENPP1 with a simple point charge (SPC) water model. Dynamic was performed with 100 ns, during the simulation the length of bond involving hydrogen was constrained using NPT ensemble, without restrains, for a simulation time of 1.2 picoseconds (ps) (temperature 300 K) was performed to relax the system.

### Cell culture

4T1 and THP-1 cells were purchased from ATCC (American Type Culture Collection) and cultured in RPMI-1640 medium (BasalMedia, L210KJ) supplemented with 10% fetal bovine serum (FBS, Meilun, PWL001) and 1% penicillin-streptomycin (PS, Meilun, MA0110) at 37°C in a 5% (v/v) CO2 atmosphere. THP-1-derived macrophages were induced by 100 ng/ml Phorbol 12- myristate 13-acetate (PMA, MedChemExpress, HY-18739) for 24 hours.

### RNA isolation, cDNA synthesis, and real-time quantitative PCR (RT-qPCR)

THP-1-derived macrophages were pretreated with various concentrations of K-756 for 30 minutes, then treated with 2 μM 2’,3’-cGAMP sodium (MedChemExpress, HY-100564A) for 12 hours. RNA-easy Isolation Reagent (Vazyme, R701-01) was used to extract total RNA from the cells, according to the manufacturer’s instructions. This total RNA was reverse transcribed into cDNA using HiScript II Q RT SuperMix (Vazyme, R223-01). RT-qPCR was conducted using ChamQ SYBR qPCR Master Mix (Vazyme, Q331-02) in the CFX96TM RealTime PCR Detection System. All the primer sequences used in this work are shown below: human ACTB forward: catgtacgttgctatccaggc, human ACTB reverse: ctccttaatgtcacgcacgat; human IFNB1 forward: cagcatctgctggttgaaga, human IFNB1 reverse: cattacctgaaggccaagga; human CXCL10 forward: ccacgtgttgagatcattgct, human CXCL10 reverse: tgcatcgattttgctcccct; human IL6 forward: ttcggtccagttgccttctc, human IL6 reverse: tacatgtctcctttctcagggc.

## Statistical analysis

Statistical analysis was performed using GraphPad Prism 9.0 software. Differences of quantitative data between groups were calculated using a 2-tailed unpaired t-test. The statistical significance level was set as *P < 0.05, **P < 0.01.

## Data availability

The data included in our paper are all from public data sets.

## Code availability

The code used to generate the results shown in this study will be available under an MIT License in the repository https://github.com/myzhengSIMM/PertKGE upon publication.

## Author contributions

S.N., X.K., Y.Z., Z.C., designed and performed the experiments, prepared the figures and wrote the manuscript; S.N. designed the PertKG and conducted the computational work; X.K. helped S.N. conduct some baseline models and computational analysis; Y.Z. contributed to the biological experiments on K-756; Z.C. and Z.W. contributed to the biological experiments on ALDH1B1 inhibitors; R.H. participated in the analysis of computational results. Z.F., X.T., N.Q., K.W., W.Z., R.Z., Z.Z., J.S., Y.W., R.Y., X.L., S.Z. and M.Z. helped check and improve the manuscript. M.Z., S.Z. and X.L. conceived, initiated, designed and supervised this study.

## Competing interests

The authors declare that they have no competing interests.

## Notes

### Competing Interest Statement

The authors have declared no competing interest.

