## Supplemental information for "Identify compound-protein interaction with knowledge graph embedding of perturbation transcriptomics"

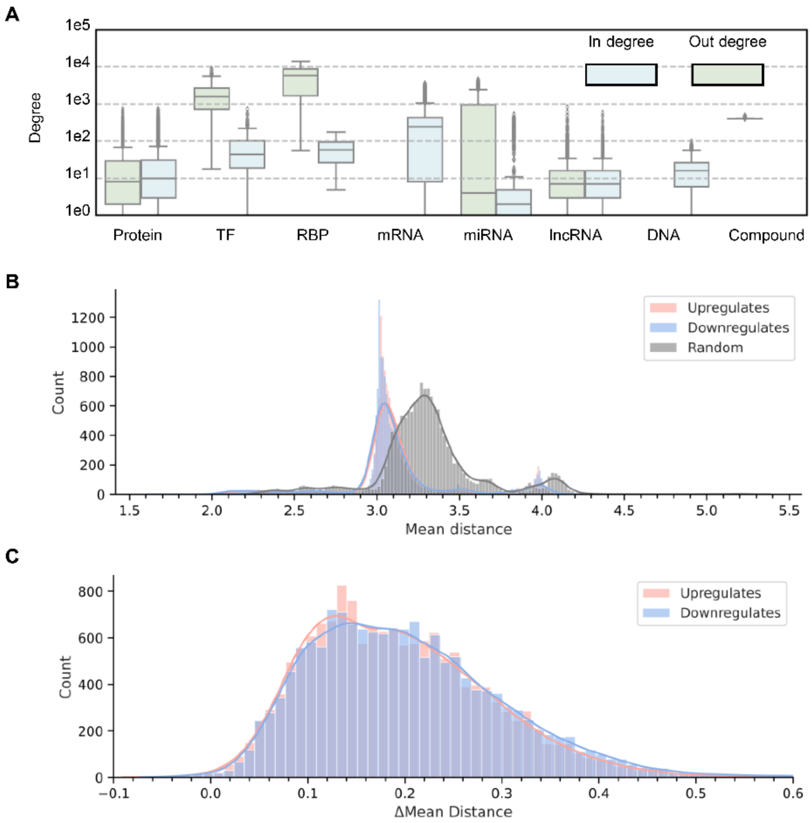


**Fig. S1. Network analysis of biologically meaningful knowledge graph. A,** Nodes' degree distribution of knowledge graph. The x-axis represents different types of nodes, while the y-axis represents the degree of nodes on a logarithmic scale. **B and C,** The shortest path analysis of differential genes. Mean distance means the average path length from DEGs to the target, while delta mean distance means the path length from random DEGs to the target substracts the path length from the observed DEGs to the target. The fundamental premise of this analysis is that, in actual biological systems' interaction networks, perturbations are more likely to lead to changes in gene expression near the target genes rather than those distant from the target.

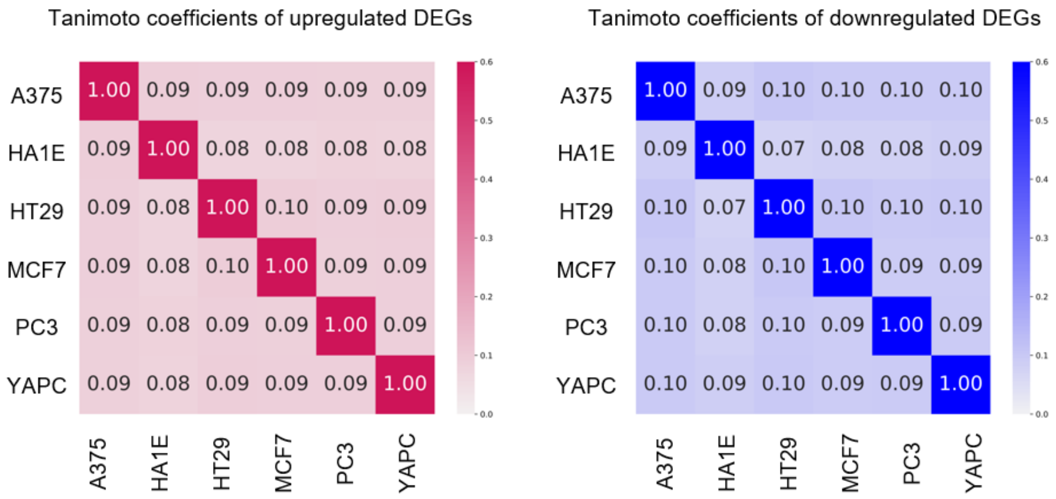


**Fig. S2. Heatmap of tanimoto coefficients of upregulated and downregulated DEGs across 6 cell lines.** The Tanimoto coefficients of upregulated and downregulated DEGs are calculated by computing a pairwise tanimoto coefficients matrix for each compound and then taking the average.

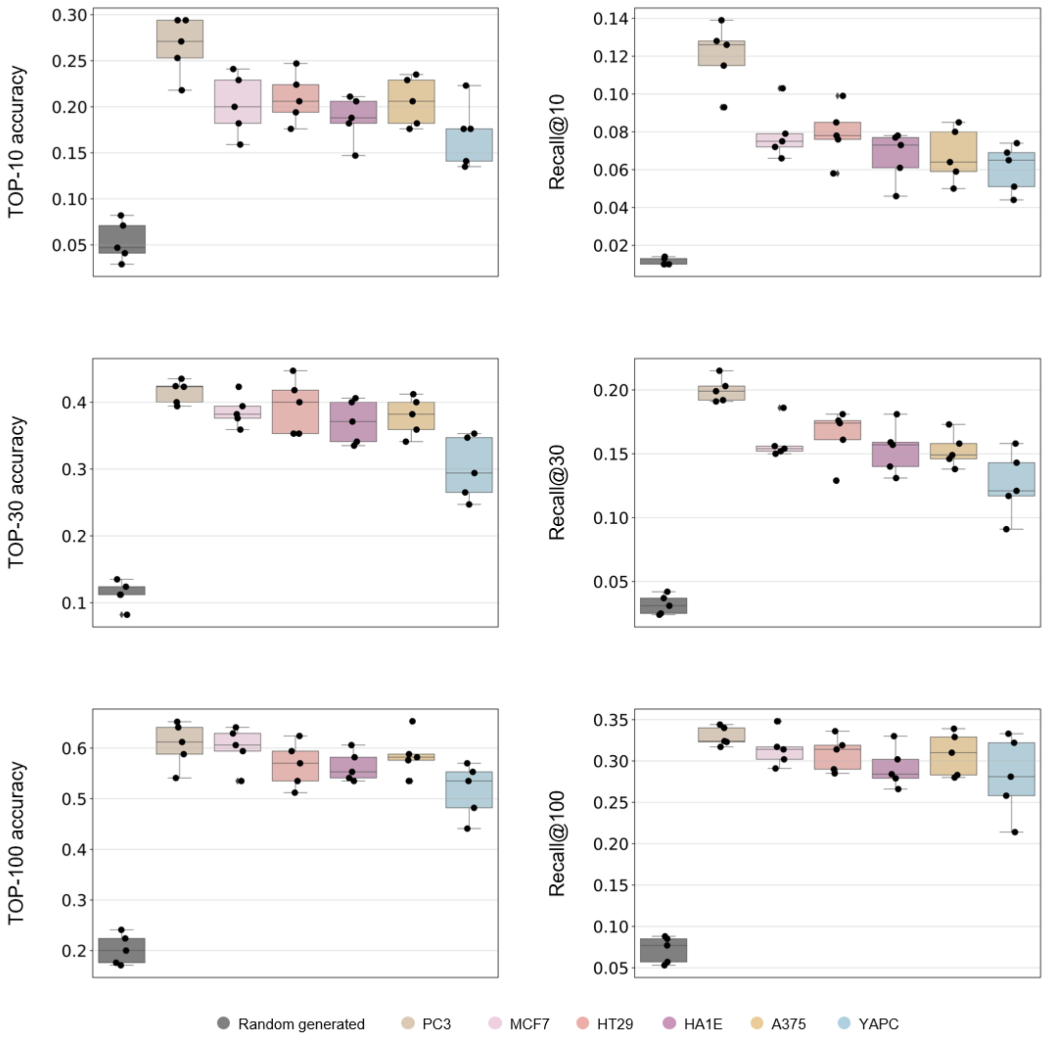


**Fig. S3. Comparison of the target inference performance for 170 compounds' DEGs in 6 cell lines and randomly generated DEGs.**

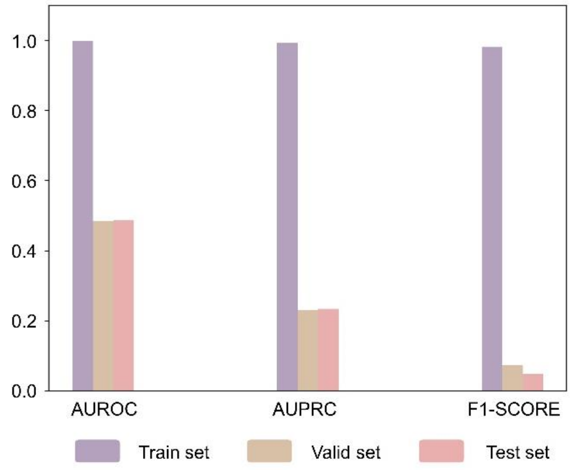


**Fig. S4. The performance of SSGCN in target cold-start setting.** To illustrate that SSGCN cannot be applied to the target cold-start scenario, we conducted a toy test. Specifically, for 6501 CPI, we randomly generated negative samples at a ratio of 1:3, and split them into train set (5,128:15,384), valid set (943:2,829), and test set (430:1,290) based on targets. While the results showed that SSGCN fits the training set, it exhibits no predictive capability on the valid and test sets.

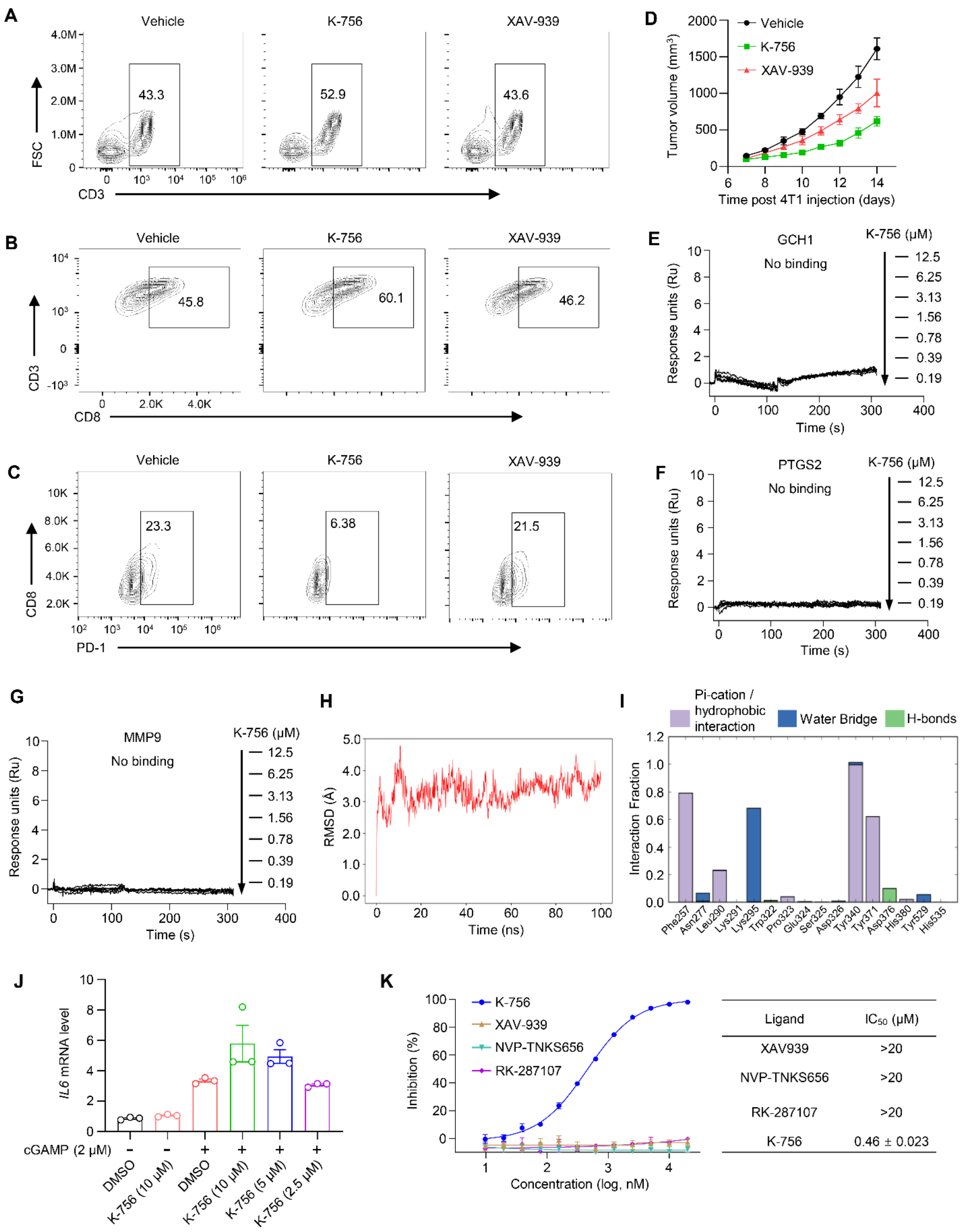


**Fig. S5. Secondary pharmacology study of K-756 by PertKGE. A-C,** Impact of K-756 and XAV-939 on the infiltration of CD3+, CD8+, and PD-1+CD8+ T cells in tumors was assessed by flow cytometry (n=4). BALB/c mice were orthotopically inoculated with 4T1 breast cancer cells and administered 30 mg/kg K-756 or XAV-939 daily via intraperitoneal injection. **D,** Tumor volumes were measured once a day. **E-G,** K-756 demonstrated no binding to GCH1, PTGS2, and MMP9 proteins in the SPR experiment. Graphs depicting equilibrium response units versus compound concentrations were plotted. **H,** Cα distance between the pocket region of ENPP1 and K-756 as a function of time in the ENPP1-K-756 simulations. **I,** The fractions of various interactions in the ENPP1-K-756 simulations. **J,** IL6 mRNA levels in THP-1-derived macrophages were measured following treatment with 2 μM cGAMP alone, or 2 μM cGAMP combined with various concentrations of K-756 for 12 h. **K,** Dose-dependent inhibition of the indicated TNKS inhibitors against ENPP1. The substrate for the ENPP1 enzymatic reaction is TMP. Error bars indicate the mean ± SEM of three biologically independent experiments (J, K)

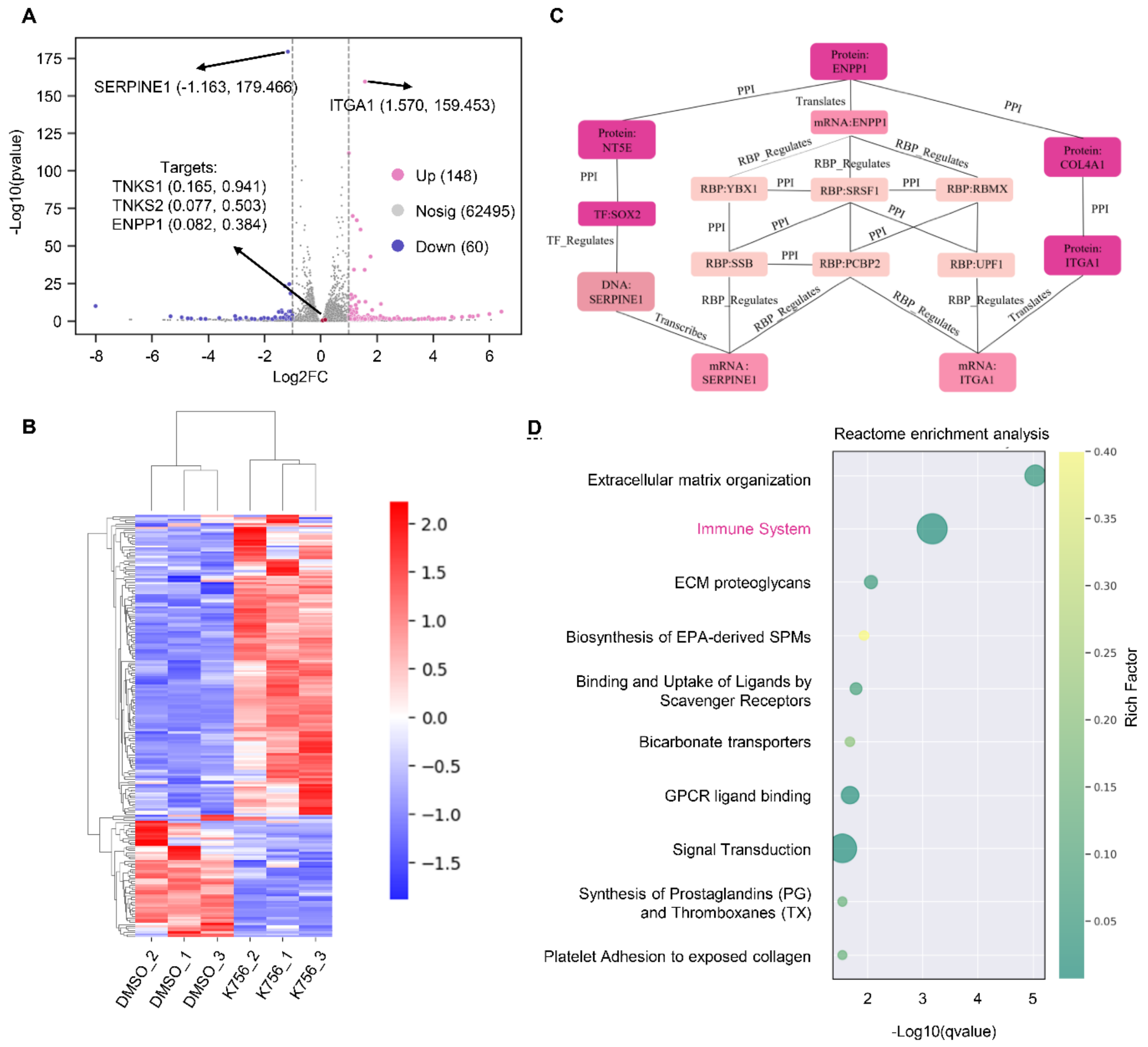


**Fig. S6. Differential analyses of K-756's profile. A,** Volcano plot to show RNA-seq results. Relative to the control group (DMSO), there are 148 significantly upregulated genes, 60 significantly downregulated genes, while the remaining 62,495 genes show no statistical significance. The expression of two known targets of K-756, TNKS1 and TNKS2, as well as our experimentally confirmed target ENPP1, only has p-values of 0.941, 0.503, and 0.384, respectively. They cannot be distinguished through differential expression analysis.**B,** Cluster analysis of all DEGs from the six samples, three belong to K756 perturbation, and the other three belong to DMSO. The color bar, ranging from blue to red, represents the low-to-high expression, respectively. **C,** Network analysis of how the DEGs are connected to the target protein ENPP1 (taking mRNA:SERPINE1 and mRNA:ITGA1, which have significant P-value, as examples) according to chemical perturbation transcriptomics-based knowledge graph. **D,** The enrichment analysis of DEGs by Reactome.

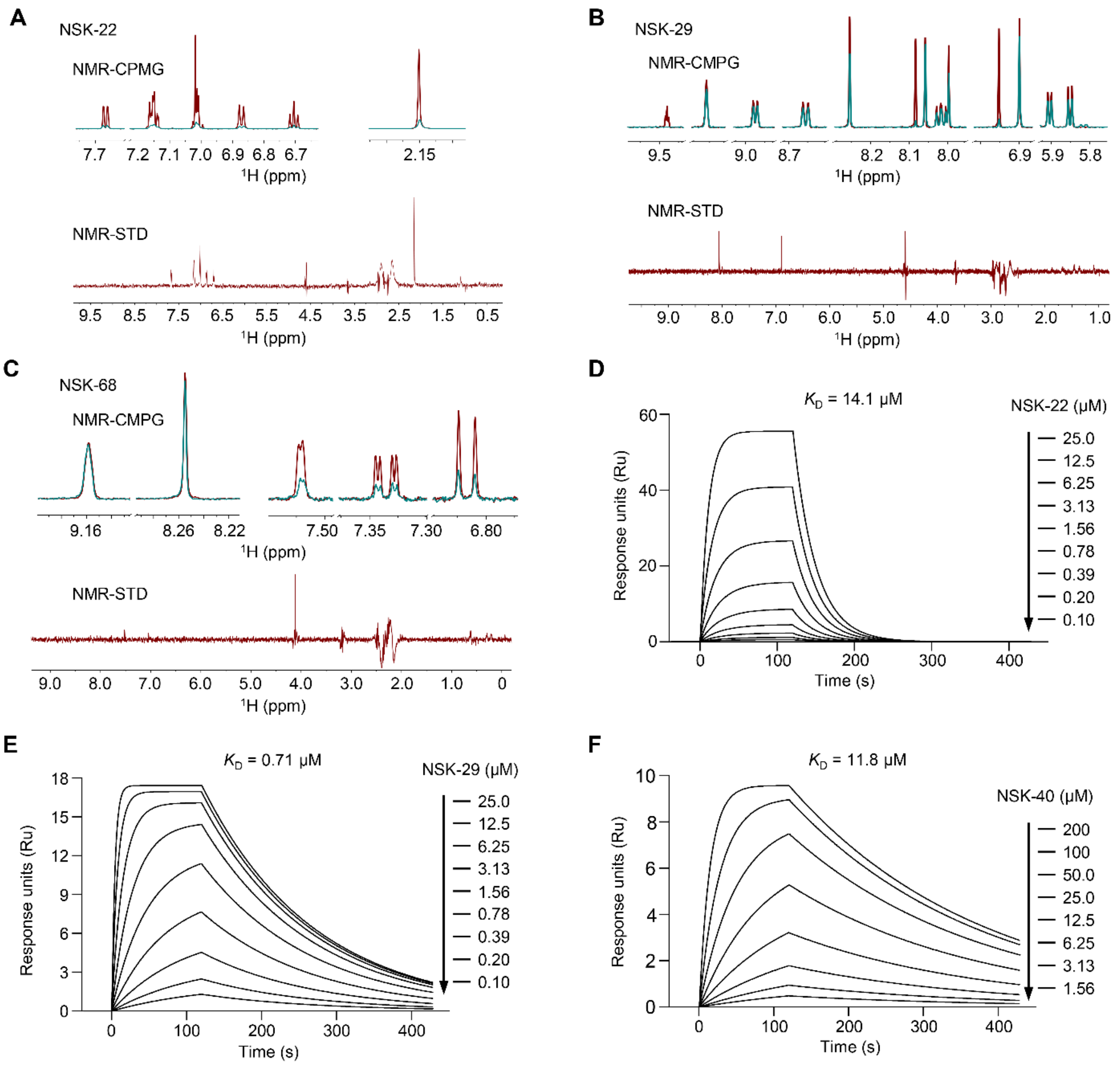


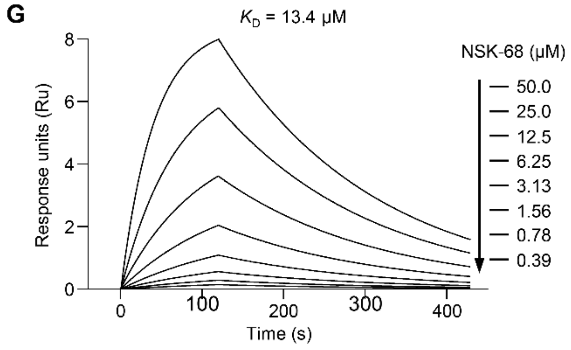


**H**

| Ligand | 7MJC |
|---|---|
| NSK-22 | 1,937 / 7,403 |
| NSK-29 | 2,145 / 7,403 |
| NSK-40 | 1,509 / 7,403 |
| NSK-68 | 6,488 / 7,403 |
| NSK-98 | 7,322 / 7,403 |

**Fig. S7. PertKGE identified five new scaffold hits for ALDH1B1. A-C,** NMR measurement of direct binding of NSK-22 (A), NSK-29 (B), and NSK-68 (C) to ALDH1B1 protein. CPMG NMR spectra for each compound (red), compound in the presence of 5 μM ALDH1B1 protein (green).

The STD spectrum for each compound was recorded in the presence of 5 μM ALDH1B1 protein. **D-G,** Binding affinity measurement of NSK-22 (D), NSK-29 (E), NSK-40 (F), and NSK-68 (G) to ALDH1B1 using SPR assay. Graphs depicting equilibrium response units versus compound concentrations were plotted. **H,** Rank of reported 5 novel ALDH1B1 inhibitors, using apo structure (PDB entry 7MJC) and Glide-SP.

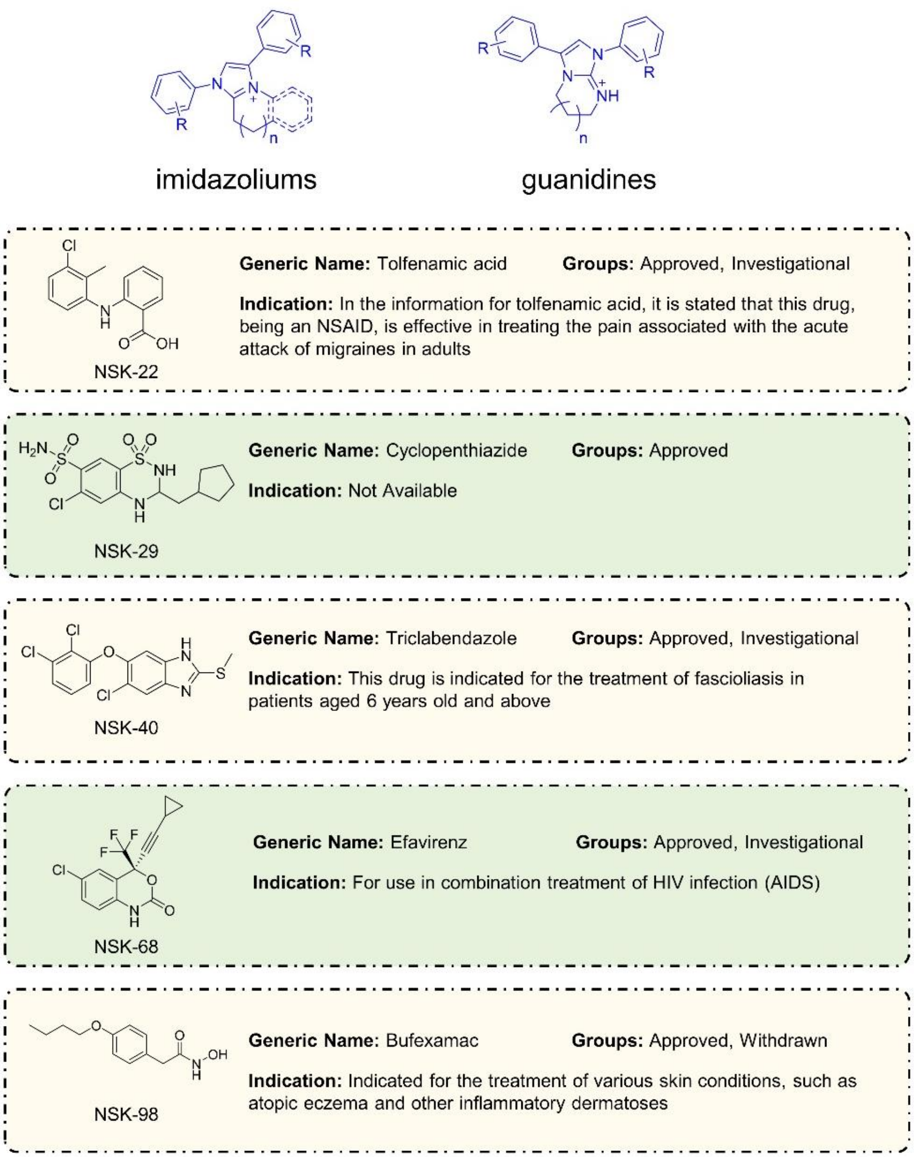


**Fig. S8. Information of ALDH1B1's inhibitors.** Pharmacology information of five hits was obtained by searching DrugBank.
